# Structural mechanism for bi-directional actin crosslinking by T-plastin

**DOI:** 10.1101/2021.12.07.471696

**Authors:** Lin Mei, Matthew J. Reynolds, Damien Garbett, Rui Gong, Tobias Meyer, Gregory M. Alushin

## Abstract

To fulfill the cytoskeleton’s diverse functions in cell mechanics and motility, actin networks with specialized architectures are built by crosslinking proteins, which bridge filaments to control micron-scale network geometry through nanoscale binding interactions via poorly defined structural mechanisms. Here, we introduce a machine-learning enabled cryo-EM pipeline for visualizing active crosslinkers, which we use to analyze human T-plastin, a member of the evolutionarily ancient plastin/fimbrin family of tandem calponin-homology domain (CHD) proteins. We define a sequential bundling mechanism which enables T-plastin to bridge filaments in both parallel and anti-parallel orientations. Our structural, biochemical, and cell biological data highlight inter-CHD linkers as key structural elements underlying flexible but stable crosslinking which are likely to be disrupted by mutations causing hereditary bone diseases. Beyond revealing how plastins are evolutionary optimized to crosslink dense actin networks with mixed polarity, our cryo-EM workflow will broadly enable analysis of the structural mechanisms underlying cytoskeletal network construction.

**One sentence summary:** Cryo-EM, biochemical, and cellular studies reveal how the crosslinking protein T-plastin bridges actin filaments in two opposing orientations.

## Main

Actin filaments (F-actin) must be incorporated into micron-scale high-order assemblies along with dozens of actin-binding proteins (ABPs) at specific subcellular locations for the cytoskeleton to fulfill its functions (*1–3*). Branched F-actin, classically associated with the plasma membrane to propel cell migration, has been extensively characterized through structural studies of the ARP2/3 complex (*4*), the sole branched-actin nucleator, culminating in a recent *in situ* sub-tomogram averaging structure with secondary-structure level detail (*5*). The other major class of actin assemblies, F-actin bundles composed primarily of co-linear filaments, remains broadly poorly understood at the protein structural level. In contrast to the central role of ARP2/3 in forming branched F-actin, F-actin bundles are compositionally and functionally diverse. They underlie the acrosome of sperm (*6*), tubular membrane protrusions including filopodia (*7*), microvilli (*8*), and stereocilia (*9*), as well as contractile networks including muscle fibers (*10, 11*), stress fibers (*12*), and the cytokinetic ring (*1*). Each type of F-actin bundle network features distinctive crosslinkers which specify its nanoscale architecture (relative filament polarities, spacings, and orientations), thereby controlling which ABPs and myosin motor proteins locally engage a network to confer its specific mechanical properties and biochemical activities. Pioneering studies of para-crystalline acrosomal bundles (*6, 13, 14*), as well as recent sub-tomogram averaging studies of muscle fibers (*15, 16*), have provided key insights into these specific network architectures, yet the detailed mechanisms of their crosslinkers remain obscure due to resolution limitations. Despite recent progress in cryo-electron microscopy (cryo-EM) analysis of individual actin filaments in complex with ABPs (*17–21*), including fragments of actin-bundling proteins (*22–25*), to our knowledge no high-resolution structures of full-length crosslinkers bridging cytoskeletal filaments have been reported, perpetuating a major gap in understanding their mechanisms.

Here we focus on human T-plastin, a member of the plastin/fimbrin family of tandem calponin-homology domain (CHD) proteins. While most crosslinkers bridge filaments through dimerization or tetramerization of subunits featuring a single actin-binding domain (ABD) (*26*), plastins contain two ABDs within a single polypeptide chain, each composed of two tandem CHDs, as well as flexibly tethered N-terminal, Ca^2+^-binding regulatory domain (RD) featuring two EF-hand motifs (Fig. 1A). When purified, fission yeast fimbrin can promote the formation of both parallel and anti-parallel F-actin bundles (*27*), suggesting individual plastin molecules must possess the capacity to bridge actin filaments in radically different geometries. Plastins are highly conserved throughout eukaryotes, suggesting they fulfill an ancient function in actin bundling (*28, 29*). Consistently, a plastin / spectrin double knockout in the *C. elegans* embryo has recently been reported to result in failure of cytokinesis (*30*), likely the most ancestral function of the actin cytoskeleton (*31*).

**Fig. 1:**
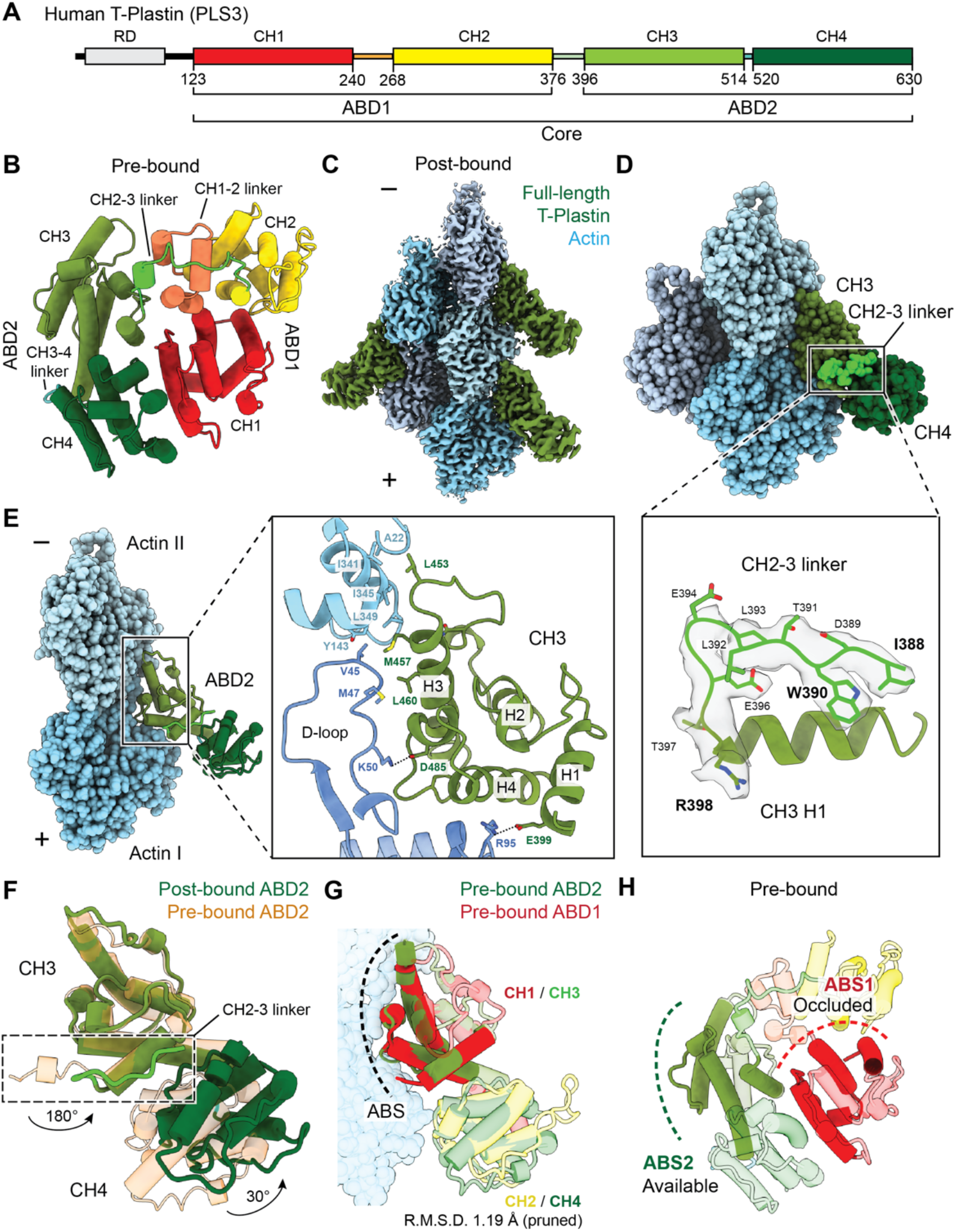
Cryo-EM structure of full-length T-plastin bound to single actin filaments resolves ABD2. (**A**) Domain structure of human T-plastin. (**B**) ‘Pre-bound’ homology model of T-plastin’s actin-binding core. (**C**) Segmented region of the ‘post-bound’ T-plastin-decorated-F-actin cryo-EM map (2.6 Å resolution) in the presence of Ca^2+^. (**D**) Post-bound T-plastin–F-actin complex atomic model. Actin subunits are displayed in varying shades of blue. CH3: olive; CH4: dark green. The inter-ABD CH2-3 linker, highlighted in bright green, is displayed along with its segmented density in the lower box. (**E**) The actin-binding interface of ABD2. (**F**) Superimposed pre-bound and post-bound models of ABD2. The conformational change of the CH2-3 linker is highlighted (box). Rotation angles indicate repositioning of the CH2-3 linker, as well as CH4 relative to CH3. (**G**) Superimposed ABD1 and ABD2 from pre-bound model on post-bound ABD2 (not shown). ABS, actin-binding site. Actin from the post-bound model is displayed. (**H**) Actin-binding sites are highlighted on the pre-bound model.

Humans feature three highly-similar plastin isoforms: I-plastin (PLS1/fimbrin), expressed in kidney, intestine, and the inner ear where it localizes to microvilli and stereocilia (*32, 33*); L-plastin (PLS2/LCP1), natively expressed in leukocytes (*34*) and ectopically expressed in many cancers (*35*); and T-plastin (PLS3), the most abundant isoform which is ubiquitously expressed in solid tissues (*36*) and dysregulated in cancers (*37*). T-plastin is a 70.8 kDa monomeric protein which functions as a key actin network stabilizer, strengthening and promoting cell protrusions by localizing to lamellipodia and filopodia in endothelial cells (*7*). Ca^2+^ suppresses its bundling activity (*38, 39*), and T-plastin is involved in many cellular processes linking calcium sensing to cytoskeletal dynamics, including cell migration (*7*), endocytosis (*27*), and membrane organization (*40*). T-plastin is associated with autosomal recessive spinal muscular atrophy (*41*) and mutations in T-plastin have recently been reported to cause congenital osteoporosis (*42–44*), likely by disrupting its Ca^2+^-regulated actin bundling activity through unclear mechanisms (*25, 45*). The actin-binding cores (lacking the flexible RD) of *S. pombe* and *A. Thaliana* fimbrin have been crystallized (*46*) in the absence of actin, and the isolated ABD2 of L-plastin (*25, 47*) and ABD1 of T-plastin (*48*) have been visualized bound to individual actin filaments with cryo-EM, providing insights into the overall structure of plastins and the actin-binding poses of single ABDs. A pioneering early electron microscopy study of negatively stained 2D para-crystalline F-actin arrays cross-linked by T-plastin provided a plausible model for the organization of a parallel bundle based on structural data available at the time (*49*), while leaving the detailed bundling mechanism undetermined.

Here, we present a general single-particle cryo-EM workflow for visualizing the structure of cytoskeletal crosslinkers actively bridging filaments, enabled by the development of a machine-learning procedure to detect and pre-sort candidate pairs of filaments with feasible 3D crosslinking geometry. We employ this pipeline, along with supporting structural, biochemical, and cell biological studies, to establish the detailed mechanism by which full-length T-plastin sequentially engages two actin filaments, allowing it to form parallel and anti-parallel bundles with nearly equivalent frequency.

### T-plastin initially engages F-actin through ABD2

While to our knowledge no crystal structures of human plastins have been solved, T-plastin’s significant homology to previously crystallized *S. pombe* / *A. Thaliana* fimbrins (*46*) (41.4% and 43.6% protein sequence identity, respectively) facilitated calculation of a reliable homology model of its two tandem ABDs (Fig. 1B, Materials and Methods), which we hereafter refer to as the ‘pre-bound’ structure (i.e. yet to bind F-actin). As reported in the crystal structures (*46*), pre-bound T-plastin adopts a closed horseshoe conformation with the N-terminal CH1 in close contact with the C-terminal CH4 (Fig. 1B). The two ABDs adopt a quasi-anti-parallel orientation, connected by the 20-residue inter-ABD CH2-3 linker (Fig. 1A,B, light green). Notably, the two intra-ABD linkers, the CH1-2 linker in ABD1, and the CH3-4 linker in ABD2, are very different in length (Fig. 1A, 28 residues and 7 residues, respectively).

We next pursued structural studies of full-length human T-plastin bound to F-actin with cryo-EM in both the absence and presence of Ca^2+^ (Fig. S1, Materials and Methods; Table S1). Inspection of micrographs confirmed that although actin filaments are bundled under both conditions, bundling is suppressed by Ca^2+^ (Fig. S1). Because the conventional Iterative Helical Real Space Reconstruction (IHRSR) approach (*50*) can only be applied to single filaments, we initially focused on the +Ca^2+^ dataset to visualize the T-plastin–single F-actin interface. We obtained a 3D reconstruction using IHRSR as implemented in RELION 3.0 (*51*) (Fig. S1; Fig. 1C) at 2.6 Å overall resolution, to our knowledge the highest resolution reported for an F-actin–ABP complex to date. Local resolutions ranged from 2.4 Å to 3.9 Å, radially decaying from the core of the filament (Fig. S2), facilitating direct atomic model building and refinement for the complete sequence of Mg-ADP α-actin and T-plastin residues 388-630 (Fig. 1D, Materials and Methods). Despite using the full-length protein, density for only 2 CHDs was observed (Fig. 1C, 1D). The map resolution allowed us to unambiguously identify them as CH3 (olive) and CH4 (dark green), i.e. ABD2 and a short segment of the CH2-3 linker (light green) at the N-terminus of CH3 (Fig.1D). The essentially uniform ABD2 decoration in the dataset indicates that when T-plastin engages F-actin, ABD2 must bind first. The robust actin binding by ABD2 also indicates that Ca^2+^ suppresses T-plastin-mediated actin bundling by inhibiting the subsequent actin binding of ABD1 after ABD2 has engaged, rather than downregulating initial F-actin binding, consistent with previous biochemical studies (*38*).

### Actin binding triggers rearrangements within ABD2, facilitating CH2-CH3 inter-ABD linker docking

Like many other ABPs (*52*), T-plastin’s CH3 domain of ABD2 engages a major site spanning the longitudinal interface of two adjacent actin protomers (Fig. 1E), which we term Actin I and Actin II (numbered from the plus (“barbed”) end of the filament), while CH4 does not directly engage actin. CHDs are composed of four major α-helices (*53*), referred to here as H1 to H4 (Fig. 1E) from N-to C-terminus, connected by loops and short, irregular helices. Helices H1, H3, and H4 constitute the Actin-Binding Surface (ABS) of CH3, mediated by an extensive hydrophobic interface between H3, Actin II, and Actin I’s D-loop, as well as salt bridges between H1/H4 and Actin I (Fig. 1E). Comparison of the actin conformation observed in a similar-resolution structure of F-actin in isolation (“bare actin”, PDB 7r8v) (*21*) versus when bound to T-plastin reveals minimal rearrangements throughout the actin structure (Fig. S2C). Compared to the pre-bound T-plastin structure, CH3 maintains its overall conformation, featuring only slight remodeling in a loop (Fig.1F), while its four major helices maintain their respective positions. However, CH4 undergoes an approximately 30° swing around the CH3-4 linker to avoid a steric clash with actin (Fig.1F), licensing a major rearrangement of the short stretch of the CH2-3 linker resolved in the cryo-EM map: a 180° degree flip away from the actin filament (Fig. 1F). As this linker connects ABD2 to ABD1, actin engagement by ABD2 is anticipated to reposition ABD1 (and potentially the RD as well).

Both CH3 and CH4 undergo minimal internal structural rearrangements when analyzed individually (Fig. S2D), suggesting that inter-domain movements through flexible linkers, rather than intra-domain rearrangements of α-helices, predominate upon T-plastin’s actin binding. The three human plastin isoforms share highly similar sequences (Fig. S3A) with at least 74% identity; consistently, our actin-bound full-length T-plastin structure is highly similar to the recently reported actin-bound structure of truncated L-plastin ABD2 (PDB 6vec, Fig. S3B) (*25*). Among all actin-bound tandem CHD ABP structures (Fig. S4A), only plastin ABD2’s second CHD could be resolved by cryo-EM (Fig. S4B-D), while the second CHDs of the utrophin, spectrin, and filamin A ABDs were all not observed, likely because plastin ABD2 has the shortest intra-ABD linker between CHDs (Fig. S4A), reducing the conformational flexibility of its actin-bound state (*54*). With the exception of plastin’s ABD2, tandem CHD ABDs engage an additional minor actin-binding site through a short N-terminal extension which folds along F-actin (Fig. S4B-D). Plastin ABD2’s unique actin binding mode likely derives from its relative position in the protein sequence: unlike other ABDs, plastin’s ABD2 is on the protein’s C-terminal end (Fig. 1A). Its preceding N-terminal element, the CH2-3 linker, engages the ABD2 itself upon actin binding and prevents it from forming an extended N-terminal actin-binding tail.

### A frustrated, pre-bundling state sterically necessitates partial disorder in ABD1

Despite modest sequence identity and the length difference between their CH1-2 and CH3-4 intra-ABD linkers (Fig. S4A), pre-bound ABD1 and ABD2 share an almost identical structure in their CHDs (Fig. 1G, RMSD 1.19 Å), indicating that when binding to F-actin, both ABDs are likely to employ a highly similar ABS (Fig. 1G), here annotated as ABS1 and ABS2 (Fig. 1H). However, the positioning of ABD1 in the pre-bound conformation is clearly incompatible with actin bundling, as ABS1 is sterically occluded within the core (Fig.1H). This suggests a stepwise bundling mechanism in which initial actin binding by ABD2 triggers a conformational change which relieves auto-inhibition of ABD1, exposing ABS1 to bind a second actin filament.

To test this hypothesis, we utilized the –Ca^2+^ dataset, where bundling is not suppressed, which we initially analyzed using IHRSR. As widespread bundling reduced the number of useable filament segments per micrograph, we obtained a map at overall lower (3.4 Å) resolution (Fig.S1; Table S1). Nevertheless, the atomic model derived from this map is indistinguishable from the +Ca^2+^ model, confirming that Ca^2+^ does not affect initial actin binding by T-plastin through ABD2 (Fig. S2A). However, unlike the +Ca^2+^ dataset, during 3D classification of the –Ca^2+^ dataset we noticed weak signal distal from ABD2 in one class, suggestive of a partially ordered additional domain (Fig. S1A, yellow circles). We therefore employed symmetry expansion to analyze each T-plastin binding site independently and extensive focused classification (Fig. S5A,B, yellow circle, Materials and Methods; Table S2) to obtain a 6.9 Å reconstruction (Fig. 2A; Fig. S5C,D) of what we term a “pre-bundling” state resolving the entire CH2-3 linker and the final three helices (H2-H4) of CH2 within ABD1. We utilized docking and a molecular-dynamics flexible fitting (MDFF) approach to build a corresponding pseudo-atomic model (Materials and Methods, Fig. 2B; Table S2).

**Fig. 2:**
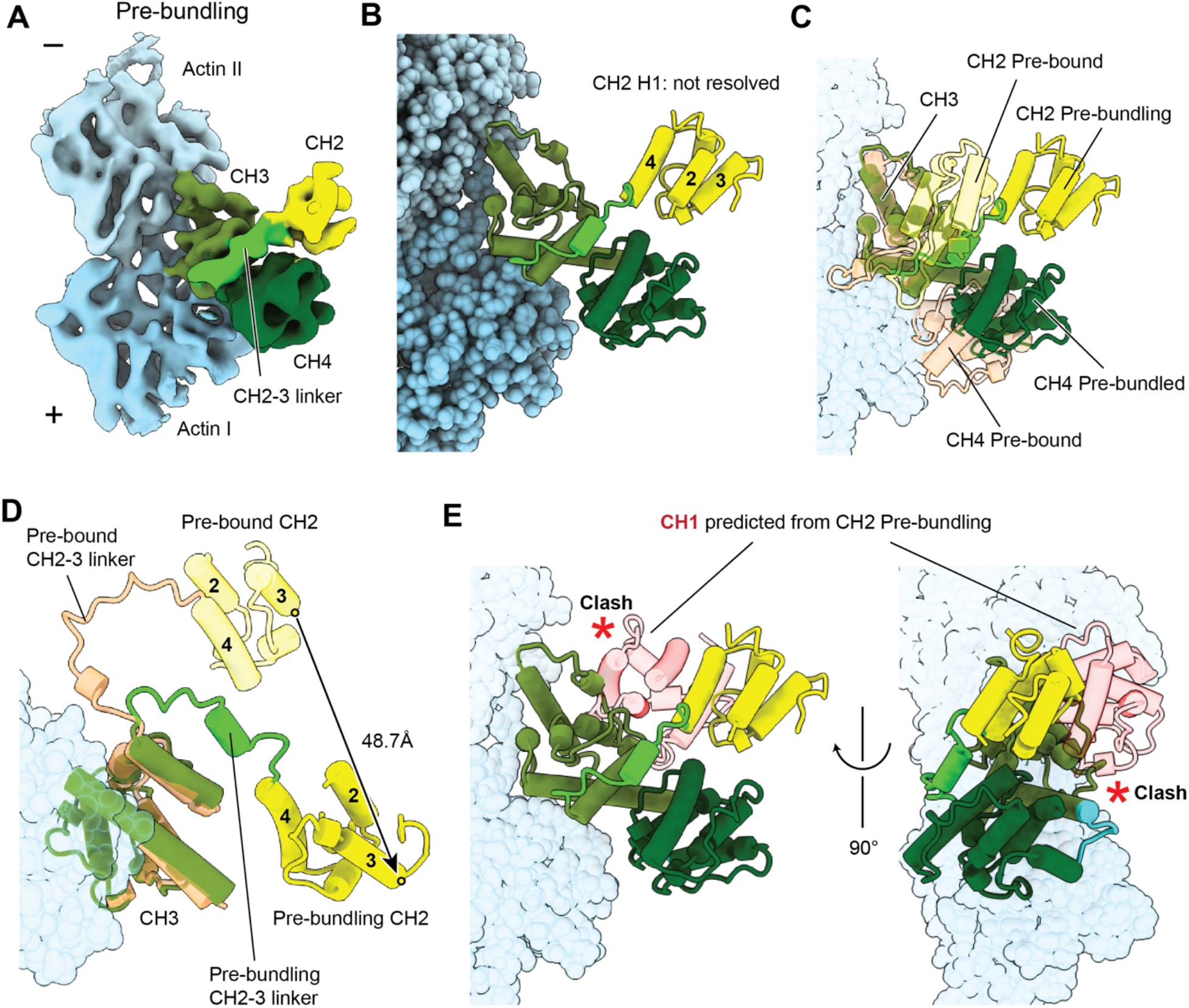
Identification of a pre-bundling conformation adopted by ABD1’s CH2. (**A**) Segmented region of the ‘pre-bundling’ T-plastin F-actin cryo-EM density map (6.9 Å resolution) in the absence of Ca^2+^ recovered by focused classification. (**B**) Flexible-fitting pseudo-atomic model of the pre-bundling state. Actin subunits are displayed in varying shades of blue. CH2: yellow; CH2-3 linker: lime; CH3: olive; CH4: dark green. (**C**) Superimposed pre-bound and pre-bundling structures of T-plastin. (**D**) Same as **C**, showing only CH2 H2-H4 and CH3. Circles represent the positions of D332’s Cα, whose displacement is displayed. (**E**) Superposition of the pre-bound ABD1 CH2 (not shown) on the pre-bundling CH2. The predicted position of CH1 results in a steric clash with ABD2, highlighted by red asterisks.

Aligning the pre-bound and pre-bundling states reveals a major conformational change within CH2 and the CH2-3 linker upon actin engagement (Fig. 2C,D): the CH2-3 linker’s approximately 120° swing results in a rotation and 48.7 Å translation of CH2 (Fig. 2D). We next examined whether this conformational transition could directly prime ABD1 for engaging a second actin filament by repositioning it as a rigid body with ABS1 available for binding. However, the CH1 position predicted by aligning ABD1 from the pre-bound state with the portion of CH2 resolved in the pre-bundling state (Fig. 2E) instead produces a major steric clash with ABD2, a result which is incompatible with a priming model. This, along with our observation that H1 of CH2 is not resolved, consistent with partial CH2 unfolding, strongly suggests the pre-bundling reconstruction represents a frustrated, metastable state where a partially disordered ABD1 can sample a large conformational space to search for a second actin filament.

### Parallel and anti-parallel bridges feature divergent ABD orientations that satisfy linker constraints

Direct structural visualization of crosslinking proteins bridging cytoskeletal filaments remains technically prohibitive, as bundles are not stoichiometrically defined protein complexes and their geometry is incompatible with current IHRSR methods (*50, 55*). To overcome these limitations, we developed a machine learning approach for specifically detecting pairs of actin filaments whose geometry is compatible with bridging by a crosslinking protein (Fig. S6, Fig.S7, Materials and Methods), the minimal unit of a bundle. Briefly, we generated synthetic datasets of paired plastin-decorated actin filaments which uniformly sample a broad range of inter-filament distances, relative orientations, and poses in projection views, which we used to train custom denoising auto-encoder and semantic segmentation neural networks. The semantic segmentation network allowed us to pick 2-filament “bundle particles” from the -Ca^2+^ dataset, detecting all the views required for 3D reconstruction (Fig. S8). Reference-free 2D class averages were analyzed by projection-matching with the plastin-decorated single filament map (Fig. S9), revealing both parallel and anti-parallel actin bundles and thereby directly confirming T-plastin crosslinks actin filaments bi-directionally. Low-resolution initial models were generated by positioning two copies of the single-filament map in 3D via joint projection-matching against 2D class averages representing nearly orthogonal views. After 3D classification, selected parallel and anti-parallel bundle particles were subsequently processed independently (Fig. S10A-C). These particles were subjected to extensive asymmetric 3D classification, uniform refinement, and finally multi-body refinement in RELION 3.1 (Fig. S10B,C; Movies S1-S2; Table S2) to obtain final sub-nanometer resolution reconstructions of both parallel and anti-parallel bundles featuring a single T-plastin bridge (Fig. S10C,D). These maps facilitated building atomic models of the ABD1-ABD2 core in both bridging conformations (Fig. 3A) utilizing a combined approach of rigid-body docking and MDFF flexible fitting (Materials and Methods; Table S2).

**Fig. 3:**
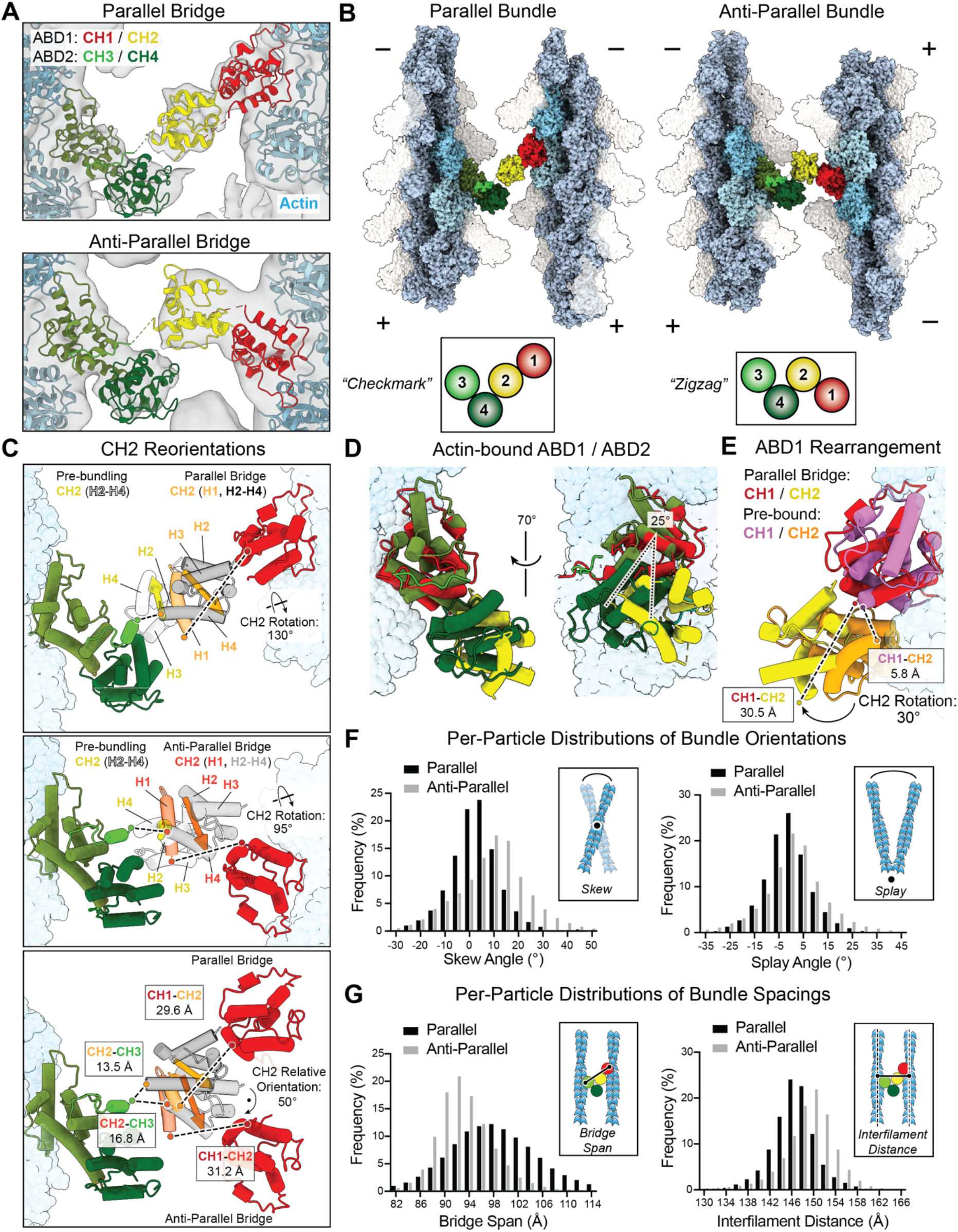
T-plastin flexibly bridges filaments in two conformations that satisfy linker constraints. (**A**) Multi-body refinement cryo-EM density maps of parallel (top, 8.2 / 8.0 Å resolution) and anti-parallel (bottom, 8.6 / 8.5 Å resolution) 2-filament bundles, with corresponding flexible-fitting pseudo-atomic models of bridging T-plastin molecules colored as indicated. (**B**) Zoomed out view of parallel and anti-parallel bundle pseudo-atomic models, highlighting their distinct T-plastin bridge conformations. (**C**) Superpositions of the pre-bundling state with the parallel bridge (top) and anti-parallel bridge (middle), as well as the parallel and anti-parallel bridges (bottom). CH2 H1 is highlighted, and CH2 re-orientations, as well as inter-CH linker lengths, are annotated. Vectors from I309 to P363 indicate the overall orientation of CH2. (**D**) Superposition of the ABD1 and the ABD2 actin-binding interfaces in the parallel bundle, aligned on actin, highlighting their distinct poses relative to the filament. (**E**) Superposition of ABD1 from the pre-bound and parallel bundle models. The extension of the CH1-2 linker (240–269) and re-orientation of CH2 relative to CH1 upon actin binding are displayed. (**F**) Per-particle distributions of indicated bundle orientation angles measured through docking analysis of multi-body refinement results. Parallel, n = 41,701; Anti-parallel, n = 28,759. (**G**) Same as **F**, but distributions of indicated bundle spacing parameters.

The structures show that in both bundle configurations, ABD1 and ABD2 bind their respective filaments in a similar fashion, with CH1/CH3 both oriented towards the filament minus ends (Fig. 3B). As a result, when bridging a parallel bundle T-plastin’s actin-binding core adopts a configuration resembling a ‘checkmark’ (Fig. 3B, left), while the anti-parallel bridge resembles a ‘zigzag’ (Fig. 3B, right). Superimposing ABD1 and ABD2 from the parallel bridge shows both ABDs bind a nearly identical site on F-actin (Fig. 3D), confirming our ABS prediction based on the high-degree of structural homology between CH1 and CH3 (Fig. 1G). However, the two ABDs protrude at angles which differ by ∼25° relative to the actin filament’s central axis (Fig.3D), consistent with a contemporary cryo-EM study of the isolated ABD1 bound to F-actin (*48*), resulting in asymmetric actin bundling by T-plastin.

We next compared the pre-bundling state to both bundled states to examine how two nearly orthogonal crosslinking geometries can be achieved (Fig. 3C, Movies S3-S4). In both cases, we observe major rotational rearrangements around the CH2-CH3 inter-ABD linker, re-positioning CH2 such that its H1 helix can re-fold, with the helix in a similar position despite the rest of the CHD adopting diverging orientations (Fig. 3C). In both cases, this produces a highly extended state of the long CH1-CH2 intra-ABD linker, which was not resolved in our reconstructions. Indeed, comparing the pre-bound and parallel bridge post-bound ABD1 conformations shows a striking rearrangement, with an ∼30° relative rotation of CH1 vs. CH2 (Fig. 3E). While this conformation enables both actin engagement by CH1 and H1 re-docking on CH2, it also requires a substantial increase in the distance between the CH1-CH2 linker’s N- and C-termini, from 5.8 Å to 30.5 Å. Although this distance can readily be spanned by the 28 AA linker, we speculate this will disrupt its compact, partially folded conformation observed in the pre-bound state (Fig.1B), consistent with it becoming flexible and unresolved in our bundle reconstructions. Superimposing CH2 from the pre-bound, pre-bundling frustrated, and bundled conformations show that, unlike the other three highly rigid T-plastin CHDs, its four major constituting helices can adopt a variety of configurations (Fig.S11A). We thus speculate that the checkmark and zig-zag conformations are the two bridging configurations which satisfy the spatial constraints imposed by the CH2-CH3 and CH1-CH2 linkers while also enabling CH2 H1 to refold, both of which can be accessed with approximately equal probability due to the structural flexibility of ABD1 in the pre-bundling state. Consistent with this model, we observe that the end-to-end distances spanned by the unresolved segments of the CH2-CH3 linker (parallel: 13.5 Å, anti-parallel: 16.8 Å) and CH1-CH2 linker (parallel: 29.6 Å, anti-parallel: 31.2 Å) are highly similar between the two bridge conformations, despite a 50° difference in CH2’s orientation (Fig. 3C, bottom).

### T-plastin forms flexible crosslinks within polarity-specific geometric constraints

As the CH1-CH2 linker and a portion of the CH2-CH3 linker were not resolved in either bundle reconstruction, and we observed substantial resolution enhancements in both reconstructions after multi-body refinement (Fig. S10D), we hypothesized both configurations would feature substantial conformational variability. To explicitly map the conformational landscape of the T-plastin bridges in our data, we implemented a high-throughput procedure to analyze the distribution of bundle geometries present. Briefly, we utilized RELION’s capacity to generate a 3D reconstruction featuring the optimal positioning of each filament detected in every bundle particle during multi-body refinement, followed by automated rigid-body docking of decorated filament atomic models whose relative positioning we subsequently analyzed (Materials and Methods). We assigned a common 3D frame of reference and defined two angles between the filaments to describe a bundle: ‘skew’, the out-of-plane tilt, and ‘splay’, the in-plane rotation (Fig. 3F). We also defined two spacing parameters: ‘bridge span’, the distance across a bridging T-plastin molecule, and ‘inter-filament distance’, the shortest distance between the central axis of each filament (Fig. 3G).

Both configurations feature broad, unimodal distributions of all 4 parameters, with no apparent correlation between skew and splay (Fig. S11B), consistent with T-plastin acting as freely flexing joint within its allowable conformation space, thereby accommodating diverse bundling geometries in the presence of mechanical perturbations. However, the conformational landscape of each configuration is distinct (Fig. 3F,G). Both the splay (mean = −1.0°; s.d. = 9.1°) and skew (mean = 2.4°; s.d. = 9.7°) distributions of the parallel bundle are centered around approximately 0°, suggesting the checkmark configuration preferentially engages collinear filaments. In contrast, while the splay angle distribution of the anti-parallel bundle is also approximately centered around 0°, (mean = 1.1°; s.d. = 12.4°), the center of its skew angle distribution is substantially displaced (mean = 9.0°; s.d. = 14.2°). These data suggest that the zig-zag conformation instead preferentially bridges filaments which deviate from collinearity. This result offers an explanation for why highly collinear actin bundles featuring plastin crosslinkers (e.g. stereocilia and microvilli) often exclusively feature parallel filaments (*9*), where many bridging plastin molecules in the checkmark conformation could accumulate along neighboring filaments to reinforce this network geometry. However, mesh-like networks (e.g. the cell cortex and lamellipodia) frequently feature mixtures of parallel and anti-parallel filaments (*56*), where our data suggest the incorporation of zigzag anti-parallel bridges would disfavor collinearity. Thus, T-plastin may preferentially adopt the checkmark or zig-zag conformation in specific subcellular contexts. Despite these distinct angular preferences, both conformations feature similar bridge spans (parallel mean 98.2 Å, anti-parallel mean: 93.1 Å) and inter-filament distances (parallel mean: 146.4 Å, anti-parallel mean: 148.9 Å), which are constrained by the dimensions of the T-plastin molecule, as predicted from previous structural analyses (*46, 49*). When the maximal width of F-actin is considered (∼70 Å), the remaining space between filaments also closely matches the width of a single filament. Our data thus suggest plastin is optimal for crosslinking dense networks, where gaps of this span should predominate (*56, 57*).

### Linker rearrangements underly T-plastin’s sequential actin bundling mechanism

Our structural data collectively suggest T-plastin employs a sequential bundling mechanism, where initial actin engagement by ABD2 triggers rearrangements in the CH2-CH3 inter-ABD linker, which docks on ABD2 to release ABD1 from an occluded, auto-inhibited state, enabling it to bind a second filament. To test this model, we designed a series of structure-guided point mutants and examined their actin binding and bundling activities. In our high-resolution actin-bound structure, CH2-CH3 linker residue W390 engages a series of hydrophobic residues along CH3 H1 (Fig. 4A), suggesting a role in mediating docking of the CH2-CH3 linker to release ABD1 after ABD2 initially engages a filament. To test this hypothesis, we generated a W390A mutant to disrupt this interface and examined its actin-binding and bundling activities via a sequential low-speed, high-speed actin co-sedimentation assay, where plastin-dependent formation of multi-filament bundles, which sediment at a lower RCF, is distinguishable from binding to individual actin filaments (*58, 59*) (Materials and Methods; Fig.4B-D). In our assay conditions (5 μM actin, 2 μM plastin), wild-type T-plastin was approximately 50% bound to actin (Fig. 4B,C, Fig. S12), and actin along with engaged T-plastin was nearly evenly divided between the low-speed and high-speed pellets (Fig. 4B,D), consistent with a mixture of bundles and individual filaments. While the overall bound fraction of the W390A mutant was indistinguishable from wild-type (Fig. 4B,C; Fig. S12), F-actin was significantly shifted to being nearly 100% in the low-speed pellet fraction, suggesting a specific effect of W390A on bundling (Fig. 4B,D), consistent with our prediction. In the pre-bound conformation, CH4 forms an extensive binding interface with CH1’s ABS (Fig. 4A), which we hypothesize stabilizes auto-inhibition of ABD1. The sequential mechanism predicts that disrupting this interface by mutating CH4 residues should enhance T-plastin’s binding and bundling activities by facilitating release of ABD1, while mutating CH1 residues should conversely suppress bundling by reducing binding to a second filament through ABD1. Consistent with this hypothesis, CH4 mutants R594A (Fig. 4B-D) and R594A R595A (Fig. S13) both bind and bundle actin significantly more that the wild-type, with both mutants and actin found almost exclusively in the low-speed pellet, while CH1 ABS mutant F191A diminishes bundling yet nevertheless strongly binds individual filaments (Fig. 4B-D), with a total actin-bound fraction indistinguishable from wild-type (Fig. S12). The R595A mutation alone, however, had minimal effect (Fig. S13), indicating that alanine substitutions in this region do not non-specifically disrupt T-plastin’s actin binding and bundling activities.

**Fig. 4:**
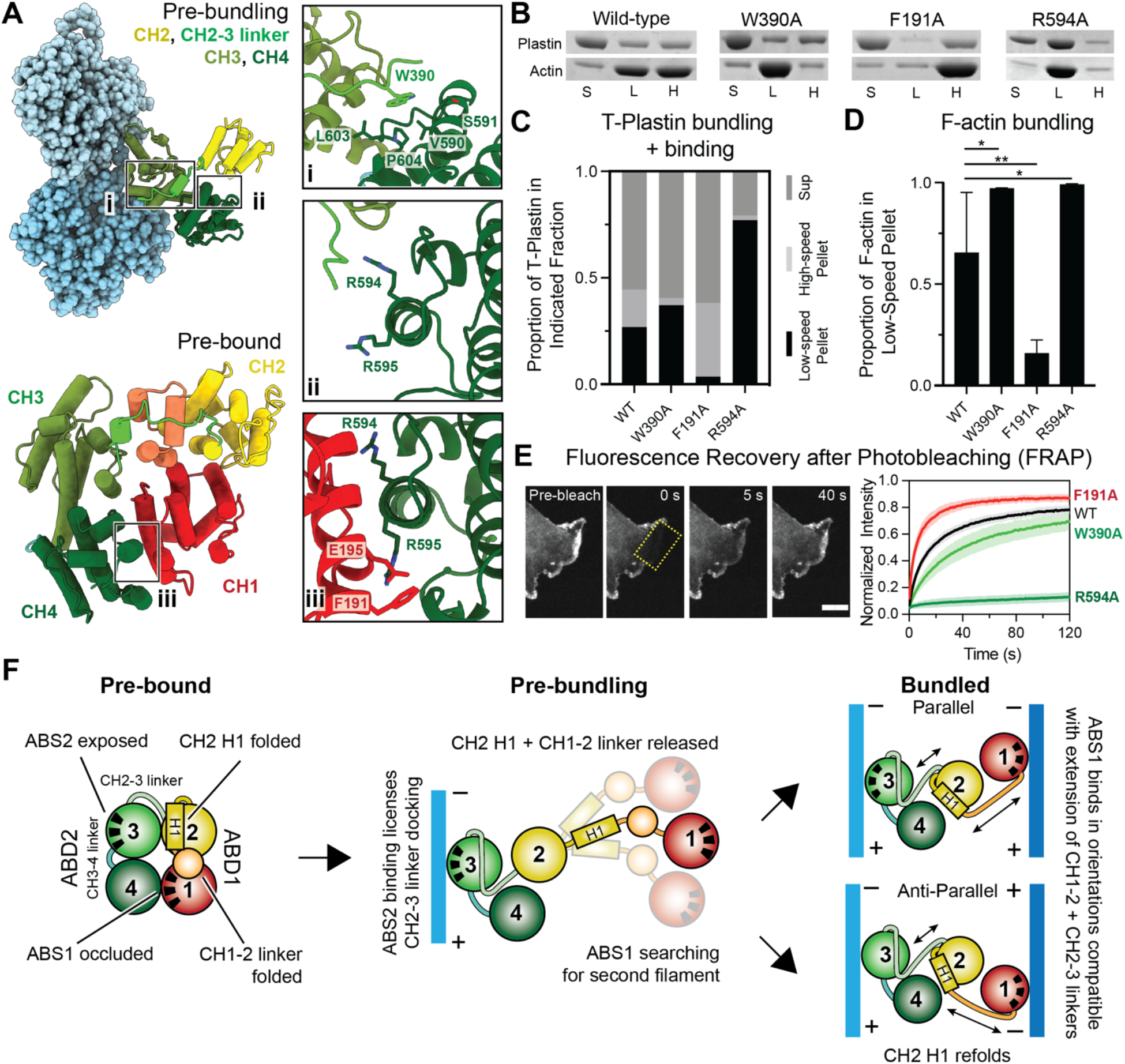
T-plastin employs a sequential mechanism to bi-directionally crosslink actin filaments. (**A**) T-plastin residues targeted for site-directed mutagenesis in the CH2-3 linker (**i**), at the pre-bound CH1-CH4 binding interface (**ii**, **iii**), and in ABS1 (**iii**). (**B**) Representative SDS-PAGE of sequential low-speed / high-speed F-actin co-sedimentation assays. S, supernatant; L, low-speed pellet; H, high-speed pellet. (**C**) Quantification of **B**: proportion of F-actin in low-speed pellet (indicative of bundling). Error bars represent S.D., 4 ≤ n ≤ 7. WT / W390A: p = 0.04; WT / F191A: p = 0.006; WT / R594A: p = 0.03. *p < 0.05, **p < 0.01, T-test. (**D**) Quantification of **B**: proportions of indicated T-plastin constructs in each fraction. 4 ≤ n ≤ 7. See Fig. S13 for statistical analysis. (**E)** FRAP assays in live HUVEC cells of the indicated eGFP tagged T-plastin constructs. Scale, bar 10 μm. (**F**) Conceptual mechanistic model.

To assess the significance of the sequential bundling mechanism in cells, GFP-tagged WT and mutant T-plastins were expressed in HUVECs to image their sub-cellular localizations and dynamics (Fig. S13A). While wild-type T-plastin and most mutants examined primarily co-localized with the ARP2/3 complex at lamellipodia, suggesting a preference for branched networks at cellular protrusions as recently reported (*7*), the enhanced bundling mutants R594A and R594A R595A exhibited a striking shift, displaying substantial localization at stress fibers and focal adhesions, where co-linear filaments predominate. A similar re-localization phenotype was reported for the phosphomimetic S406E L-plastin mutant, which is also anticipated to disrupt inter-ABD association (*48*). Collectively, these data suggest that plastin auto-inhibition and the sequential bundling mechanism is critical for actin network selection, thereby governing sub-cellular localization. We next pursued fluorescence recovery after photobleaching (FRAP) assays in cells where F-actin assembly and disassembly were pharmacologically arrested, facilitating specific probing of plastin binding dynamics (Materials and Methods, Fig. 4E, Fig S13). In these studies, we interpret rapid recovery to broadly correspond with low bundling activity, indicating T-plastin is readily displaced from actin networks *in vivo*, while slow recovery is conversely indicative of strong bundling activity (*7*). Consistent with our *in vitro* binding assays, W390A showed significantly slower recovery than the wild-type (Fig. 4E), with a half-time (t_1/2_) of 34.6 versus 17.7 s (Table S3), indicative of enhanced bundling. Conversely, F191A displayed more rapid recovery (Fig. 4E; t_1/2_ 8.8 s, Table S3), indicative of weaker bundling. The R594A (Fig. 4E; t_1/2_ 212.9 s, ∼75% immobile, Table S3) and R594A / R595A mutants (Fig. S13) both showed minimal recovery, indicative of extremely stable engagement of actin networks, while the R595A mutant alone once again had minor effects (Fig. S13). In summary, the sequential mechanism broadly predicted the impacts of T-plastin mutations both upregulating and downregulating bundling *in vitro* and in cells.

To assess whether the model could also rationalize osteoporosis-linked T-plastin mutations reported in human patients, we mapped these mutations onto our pre-bundling and post-bundling structures (Fig. S11C). Other than two mutations in CH3, N446S and L478P, that likely impact the ABD2–actin binding interface (*25*), all other mutations cluster at the center of the T-plastin bridge in the regions coordinating the structural transitions we describe here, including interfaces between CH4 and CH2, and the CH1-CH2 linker, suggesting their pathophysiology is likely linked to disruption of the conformational transitions required for bi-directional F-actin bridging.

## Discussion

Here, we introduce a general method for visualizing the basic structural unit of cytoskeletal networks with cryo-EM, filaments bridged by crosslinking proteins, which we have utilized to establish how the evolutionarily-ancient family of plastin/fimbrin tandem CHD proteins bi-directionally crosslink actin filaments. Our structural, biochemical, and cell biological data support a sequential actin bundling mechanism (Fig. 4F), in which structural transitions in the flexible linkers between T-plastin’s 4 CHDs play a predominate role in the conformational contortions that enable the protein to stably engage two nearly oppositely oriented filament geometries. We speculate that the formation of the meta-stable, single-filament-engaged pre-bundling state plays a key role in enabling ABD1 to conduct a broad search for a second filament. This can resolve into two stable bridging conformations, the parallel ‘checkmark’ and anti-parallel ‘zigzag’, likely due to a complex balance of forces exerted through linker dynamics, CH2 H1 folding, and ABD1 actin binding. While both ABDs engaging actin will stabilize a bridge, CH2 H1 refolding requires a CH2 orientation that substantially extends the long CH1-CH2 linker, which is likely unfavorable versus its folded configuration in the pre-bound state (Fig. 4F). Thus, while our multi-body analysis clearly supports substantial flexibility in both plastin bridge conformations (Fig. 3F,G), we speculate the CH1-CH2 linker acts as an internal spring-like element which restricts their conformational landscapes. While our studies do not directly inform upon the mechanism of Ca^2+^ regulation due to the RD remaining unresolved in all of our structures, the sequential mechanism presented here is fully compatible with a previously proposed model in which the RD engages and downregulates the actin-binding of ABD1 in the presence of Ca^2+^ to suppress bundling (*39*).

We provide detailed structural insights into how plastins serve as versatile crosslinkers of dense actin networks with multiple filament polarity organizations, likely an important feature for an ancestral crosslinker family required to broadly fulfill actin crosslinking functions in ancient eukaryotes. The diversification of crosslinkers featuring CHD actin-binding domains, which form well-defined homomers with stricter geometric requirements (e.g. α-actinins, spectrins, filamins, etc.) (*53*), thus likely tracked with the capacity of complex cells to build sub-cellular actin networks with specialized properties. Our analysis highlights how nanoscale structural transitions in a crosslinking protein can dramatically alter the mesoscale geometric properties of the actin network it builds, and other crosslinkers may also populate complex structural landscapes which impact cytoskeletal self-organization. The approach we introduce here should broadly enable high-resolution structural analysis of active crosslinking proteins to reveal the mechanistic principles underlying cytoskeletal network construction, as it is readily extensible to diverse multi-filament assemblies.

## Acknowledgements

We gratefully acknowledge Mark Ebrahim, Johanna Sotiris, and Honkit Ng from the Rockefeller University Cryo-EM Resource Center (CEMRC). Additional cryo-EM data collection was conducted at the Simons Electron Microscopy Center (SEMC) and the National Resource for Automated Molecular Microscopy (NRAMM) located at the New York Structural Biology Center.

## Author contributions

**L.M.:** Conceptualization, Methodology, Investigation, Formal analysis, Visualization, Writing - Original draft. **M.J.R.:** Conceptualization, Methodology, Investigation, Formal analysis, Visualization, Writing - Original draft. **D.G.:** Methodology, Investigation, Formal analysis, Visualization. **R.G.:** Methodology, Investigation. **T.M.:** Conceptualization, Formal analysis, Project administration, Funding acquisition, Supervision. **G.M.A.:** Conceptualization, Formal analysis, Visualization, Project administration, Funding acquisition, Writing - Original draft, Supervision. **All authors:** Writing – Review & editing.

## Data and materials availability

The atomic coordinates for T-plastin–F-actin complexes have been deposited in the Protein Data Bank (PDB) with accession codes: 7R94, high-resolution “post-bound” model; 7SXA, pre-bundling model; 7SX8, parallel bundle model; and 7SX9, anti-parallel bundle model. Cryo-EM density maps have been deposited in the Electron Microscopy Data Bank (EMDB) with accession codes: EMD-24323, 2.6 Å high-resolution map (+Ca^2+^); EMD-25496, 3.4 Å high-resolution map (–Ca^2+^); EMD-25496,pre-bundling map; EMD-25494, parallel bundle map; and EMD-25495, anti-parallel bundle map. Custom software is available at https://github.com/alushinlab/plastin_bundles as open source. The primary datasets generated during this study contributing to Fig. 1-4 and Fig. S1-S13 are available from the corresponding author (G.M.A.) upon request without restriction.

---

**Fig. S1:**
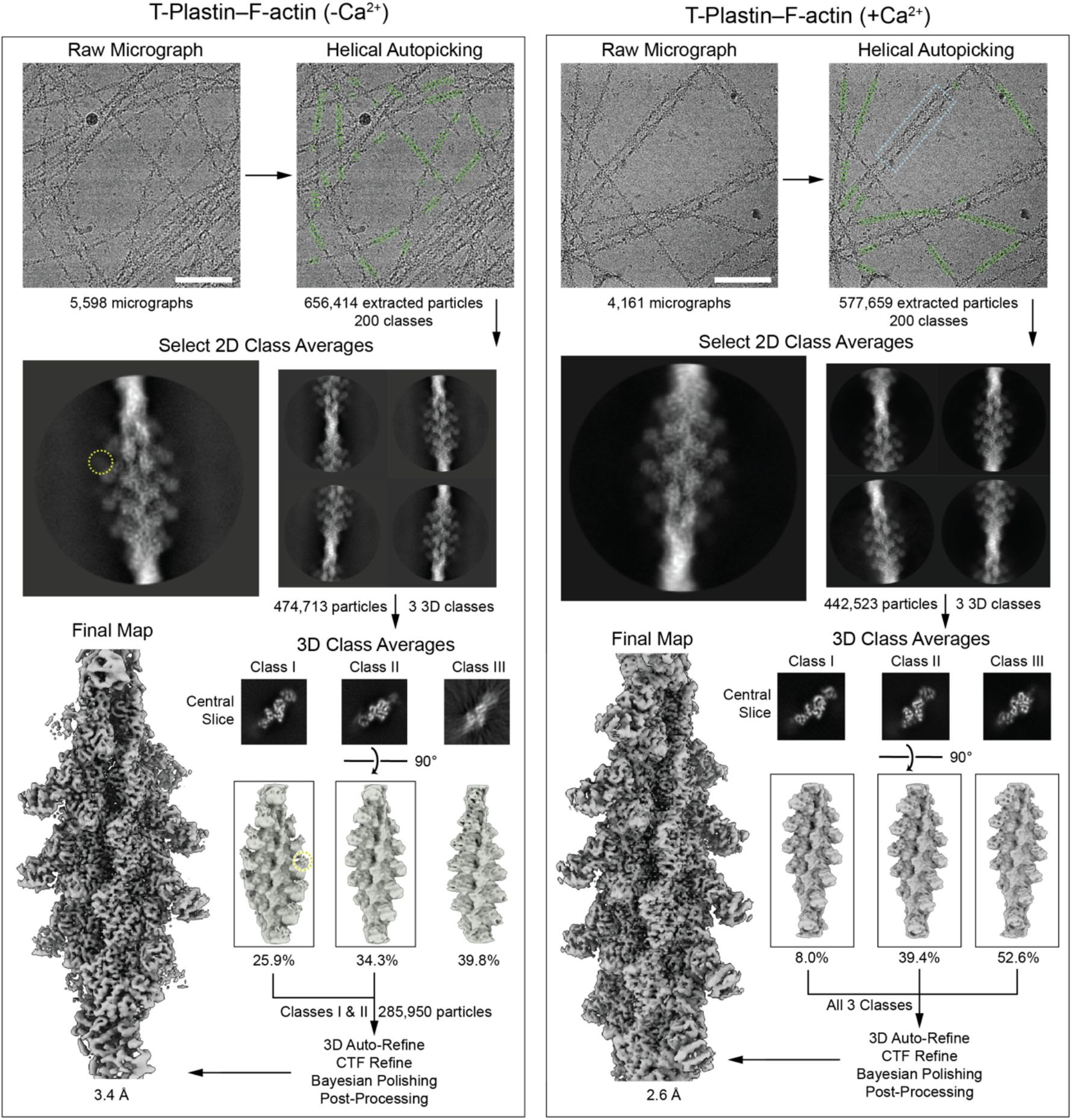
Cryo-EM data processing workflow for single-filament T-plastin–F-actin complexes. Single-filament single particle helical cryo-EM processing workflow for T-plastin decorated F-actin datasets in the absence (left) and presence (right) of calcium. Scale bars, 100 nm. Yellow circles: weak signals in 2D and 3D classes of the -Ca^2+^ dataset targeted for subsequent focused classification. Blue box: a representative 2-filament F-actin bundle.

**Fig. S2:**
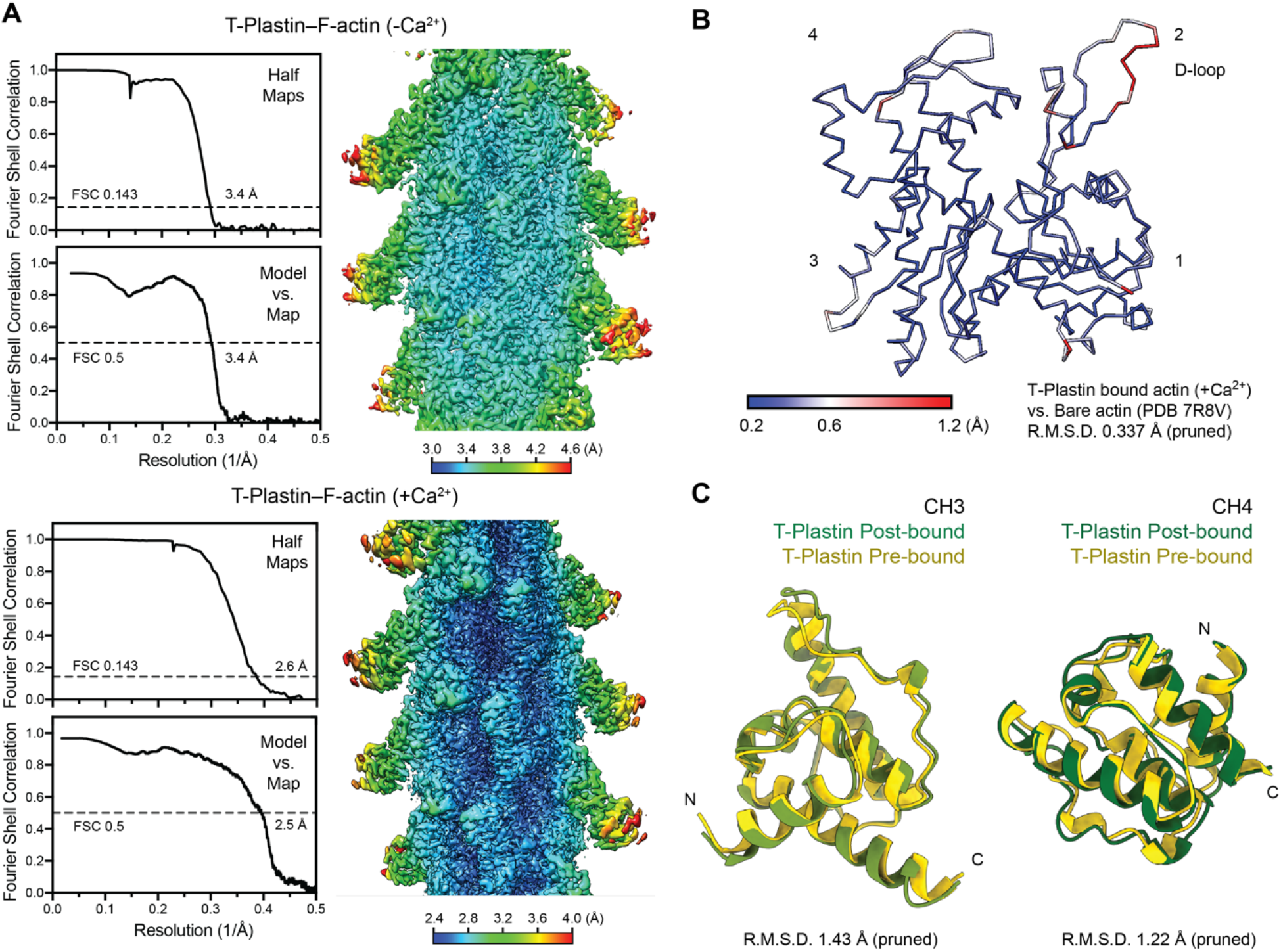
Resolution assessment and analysis of single-filament T-plastin–F-actin complexes. (**A**) 3D reconstruction and map-model analyses (half-map Fourier Shell Correlation (FSC), map-to-model FSC, and local resolution) for datasets in the absence (top) and the presence (bottom) of calcium. (**B**) Actin protomer in Cα representation, colored by per-residue RMSD between the T-plastin–F-actin +Ca^2+^ structure (this work) and bare F-actin (PDB 7r8v). Numerals indicate actin subdomains. (**C**) Individual superpositions of the pre-bound and post-bound CH3 and CH4 domains.

**Fig. S3:**
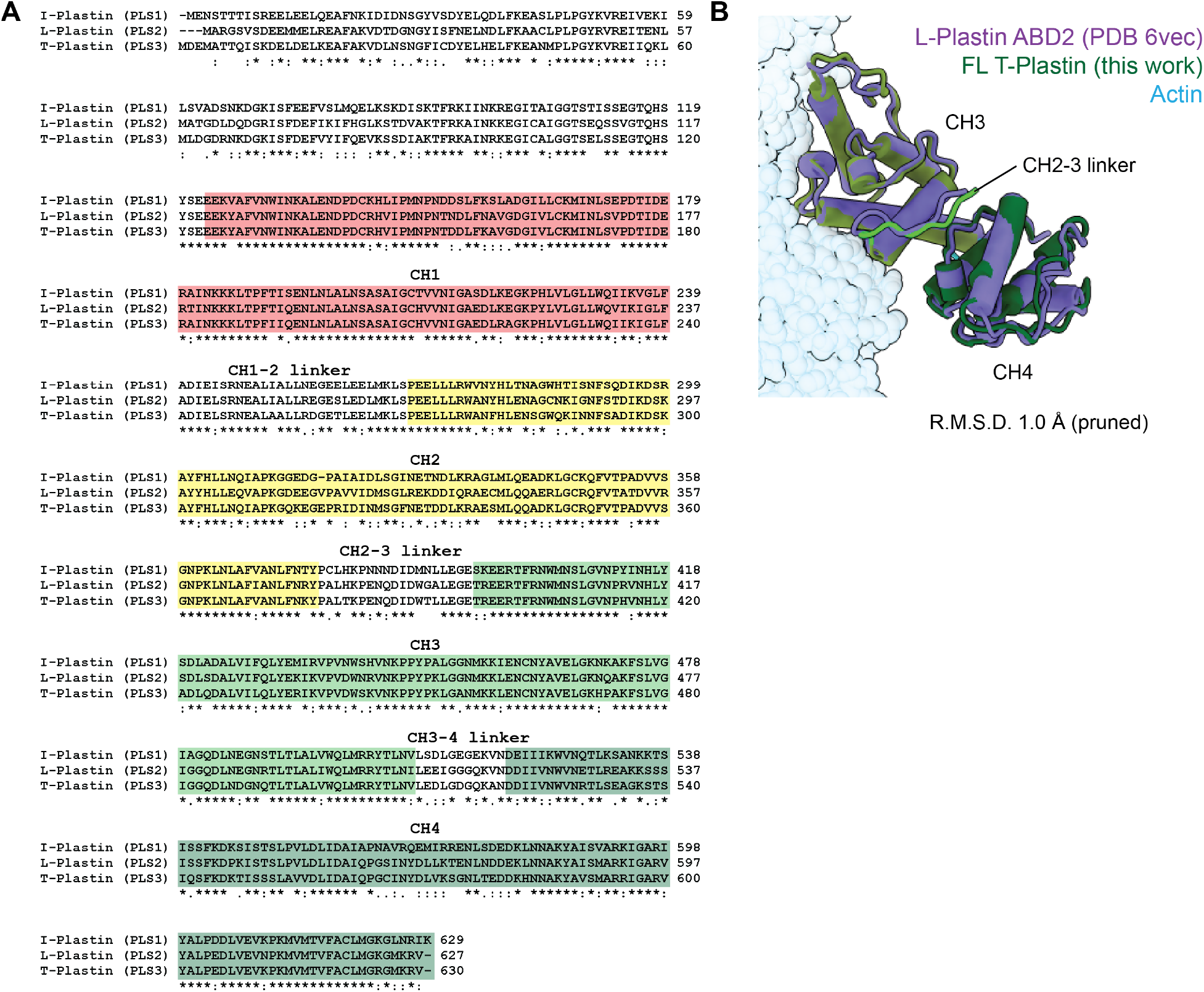
Multiple sequence alignments and structural comparison of human plastin isoforms. **(A)** Multiple sequence alignment of human I-plastin (PLS1), L-plastin (LCP1/PLS2), and T-plastin (PLS3). **(B)** Superposition of F-actin bound L-Plastin ABD2 (PDB 6vec) and full-length T-plastin (this work).

**Fig. S4:**
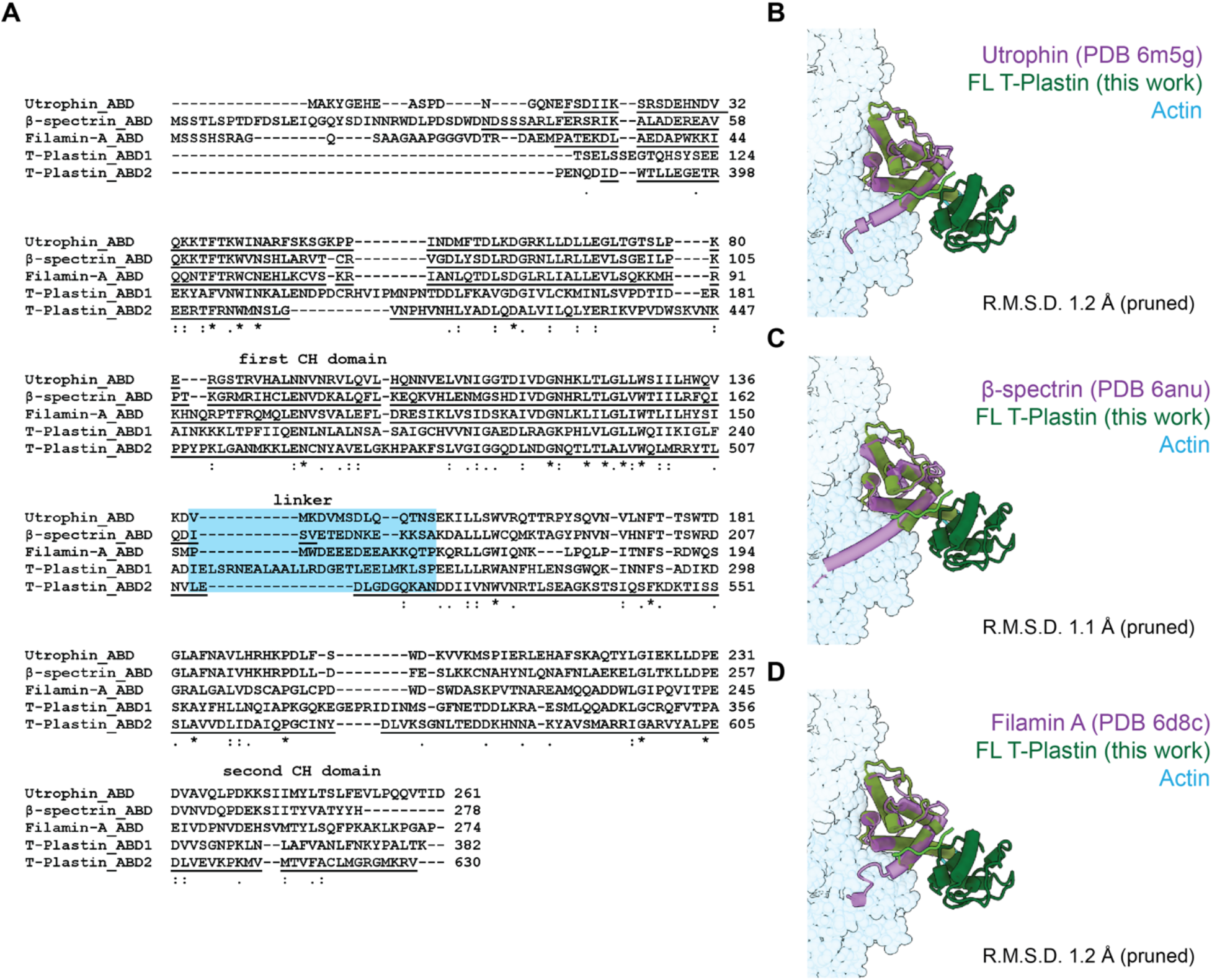
Multiple sequence alignments and structural comparisons of actin-binding CH domains. (**A**) Multiple sequence alignment of the ABDs of human utrophin, β-spectrin, filamin A, and T-plastin. (**B-D**) Superpositions of F-actin bound utrophin ABD (PDB 6m5g, **B**), β-spectrin ABD (PDB 6anu, **C**), or filamin A ABD (PDB 6d8c, **D**) with T-plastin ABD2 (this work).

**Fig. S5:**
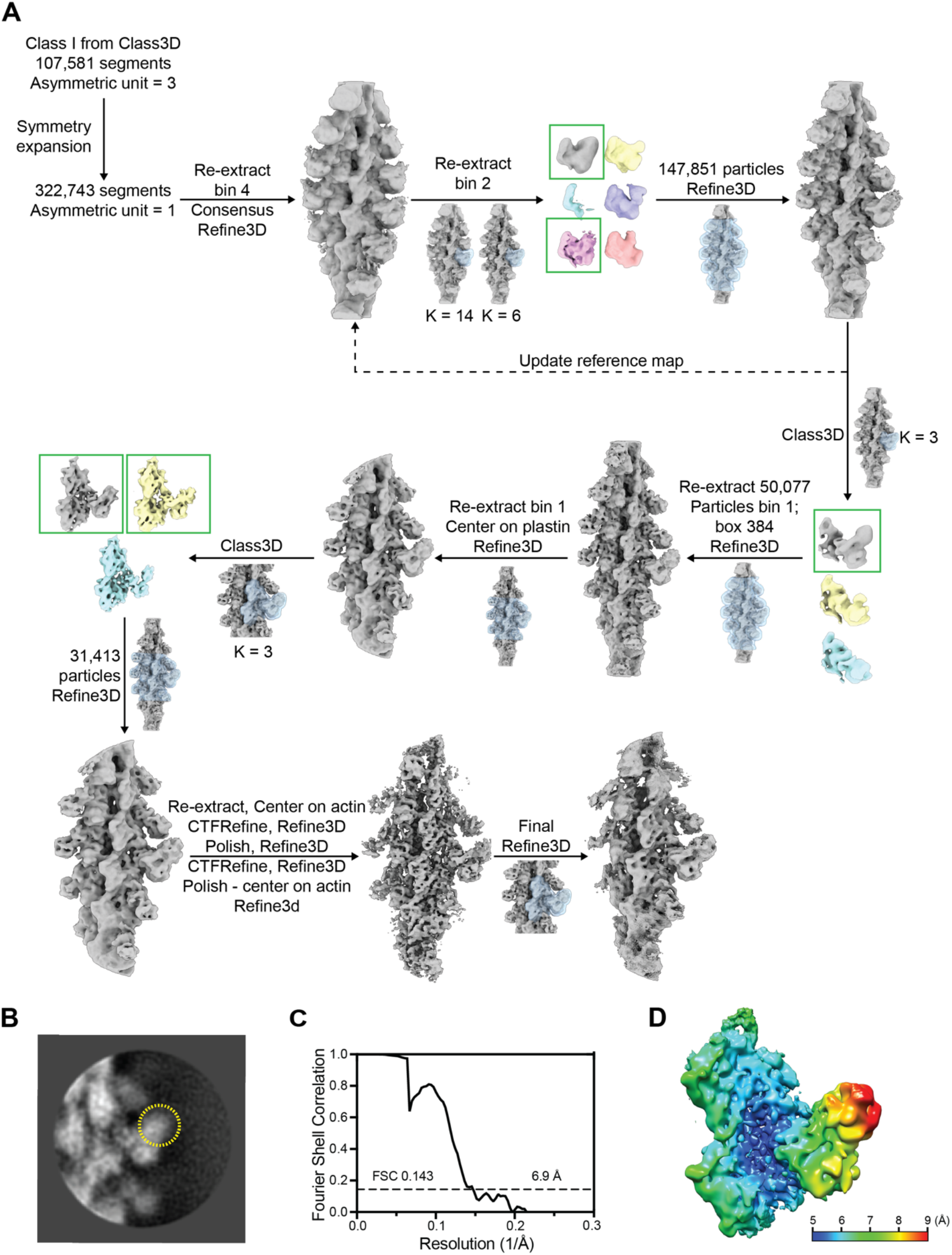
Cryo-EM data processing workflow and data analysis for the pre-bundling state. (**A**) Cryo-EM data processing / focused classification workflow to visualize the pre-bundling state of the T-plastin–F-actin complex in the absence of calcium. (**B**) Representative 2D class average after masked 3D classification. Signal beyond ABD2 is highlighted in the yellow circle. (**C**) Half-map Fourier Shell Correlation (FSC) analysis of the final reconstruction. (**D**) Local resolution analysis of the final reconstruction.

**Fig. S6:**
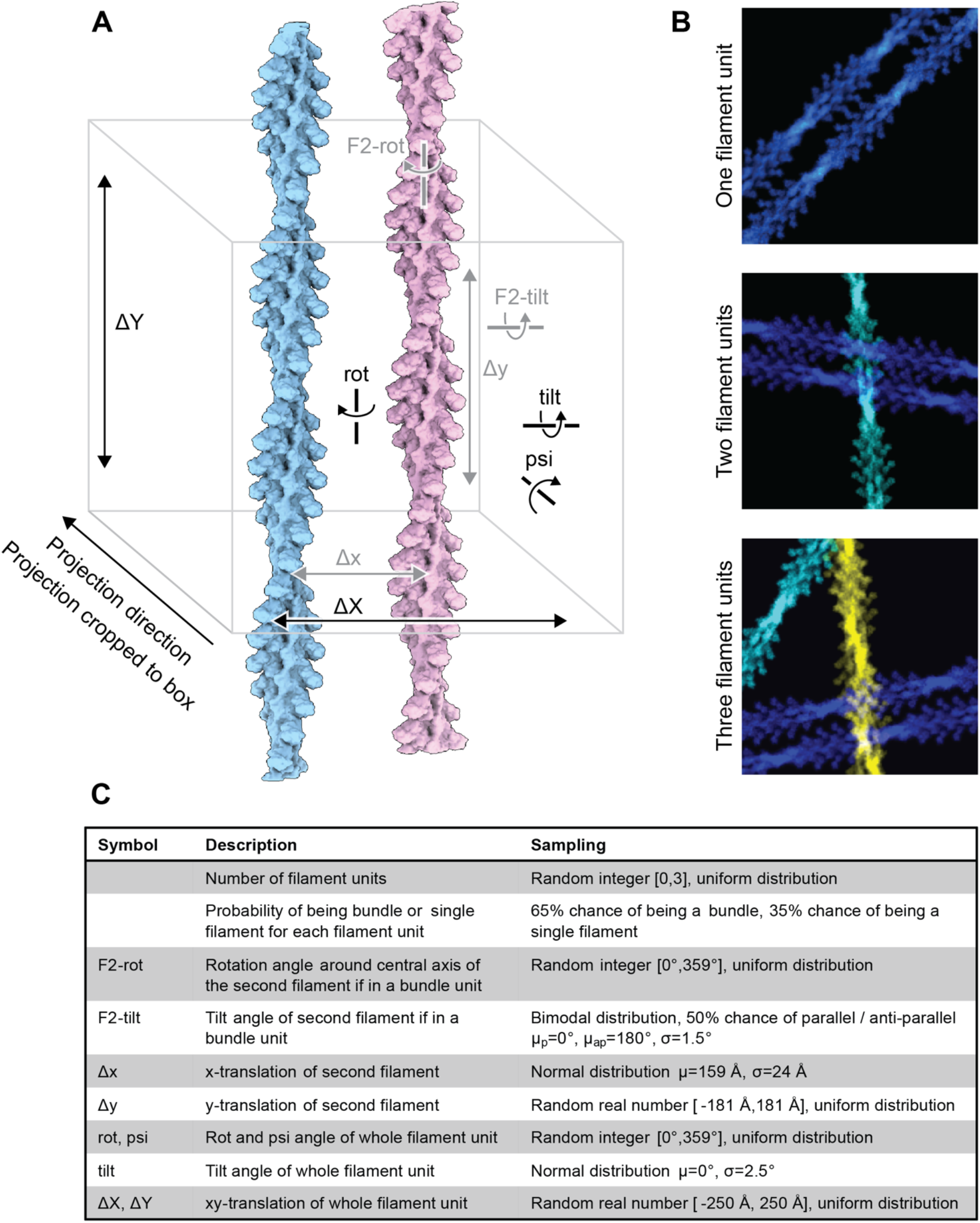
Generation of synthetic multi-bundle images for neural network training. (**A**) Diagram of 3D volume transformations applied to two T-plastin decorated filaments to generate plausible projection images of crosslinked actin filaments. (**B**) Pseudo-colored example synthetic projections with one (top), two (middle), and three (bottom) filament units. (**C**) Summary of parameter space sampled to generate synthetic datasets of plausible particle images of varying filament and bundle number.

**Fig. S7:**
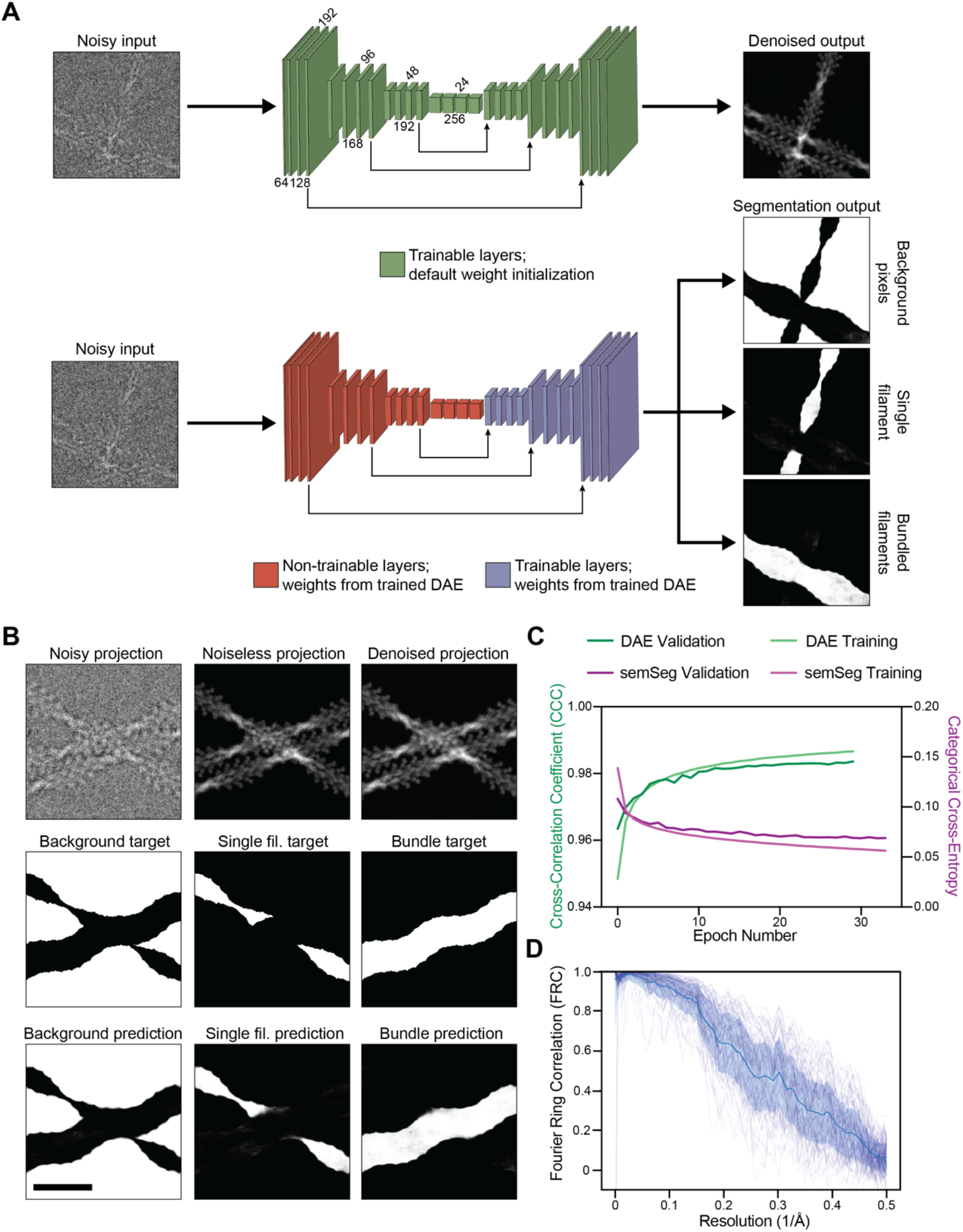
Bundle picking neural network architecture and performance on synthetic datasets. (**A**) Schematic of data processing workflow for denoising autoencoder (top) and fully convolutional neural network for semantic segmentation (bottom). (**B**) Representative denoising and semantic segmentation performance on synthetic particle image containing one 2-filament bundle and one single filament. Scale bar, 30 nm. (**C**) Neural network training metrics across epochs. DAE: denoising auto-encoder; SemSeg: semantic segmentation. (**D**) Noiseless projection (ground truth)-to-denoised projection Fourier Ring Correlation (FRC) curves from 100 example images. Mean FRC +/- 1 S.D. computed from 10,000 randomly sampled images is shown in dark blue.

**Fig. S8:**
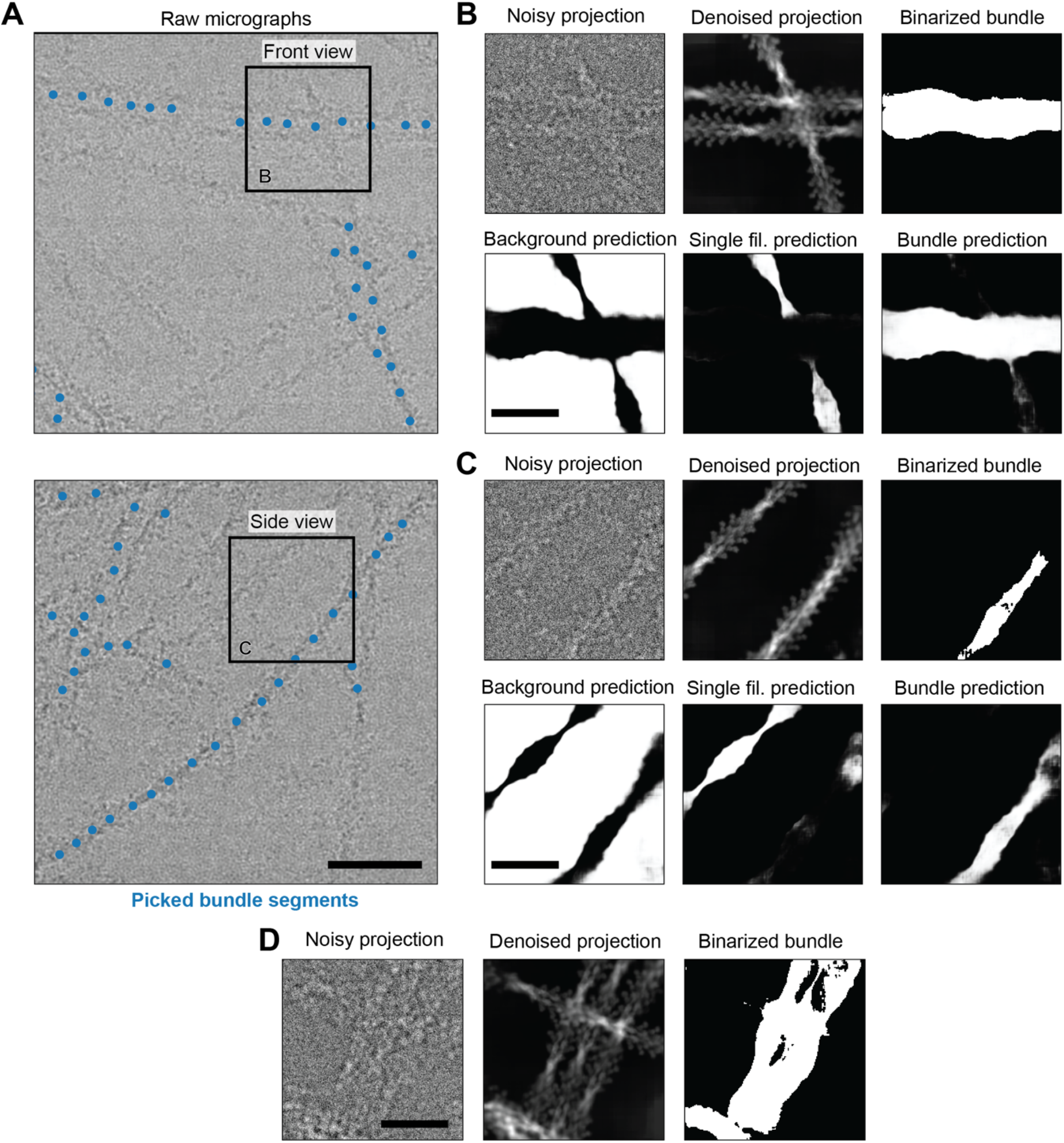
Neural network performance on experimental cryo-EM data. (**A**) Example micrographs featuring single filaments and bundles, highlighting “railroad track” front view (top) and side view (bottom) of bundled filaments. Scale bar, 60 nm. (**B**) Network performance on front view particle from **A**. Top row, left: extracted particle image; middle: denoised particle image; right: binarized bundle channel used for particle picking. Bottom row, semantic segmentation channels denoting background, single filaments, and bundle. Scale bar, 30 nm. (**C**) Network performance on side view from **A**, which is not readily discriminated from single filaments by eye. Subpanels as in **B**, scale bar 30 nm. (**D**) Picked particle featuring a three-filament bundle, upon which the network was not explicitly trained. Scale bar, 30 nm.

**Fig. S9:**
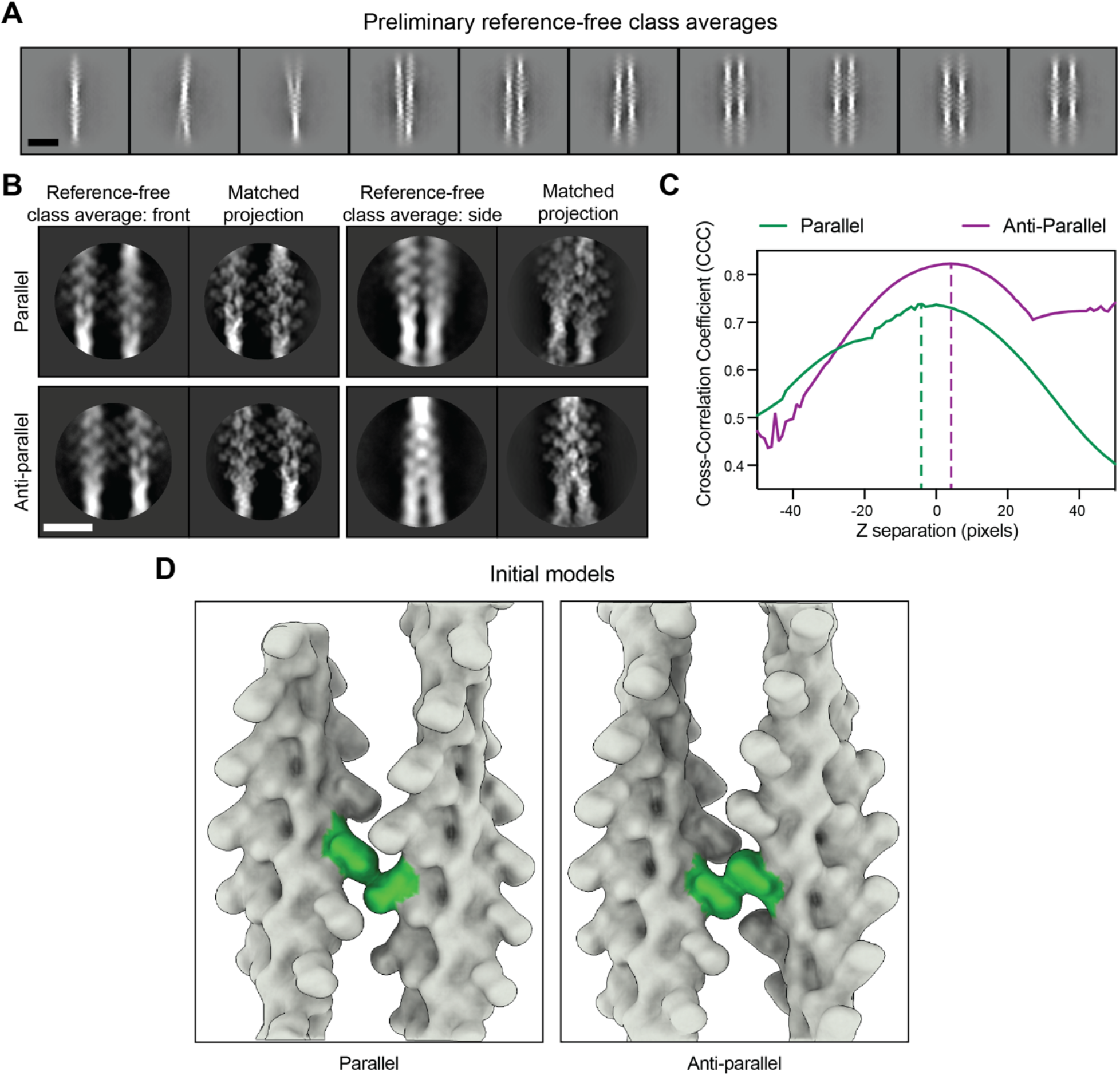
Bundle cryo-EM 2D analysis and initial 3D model generation. (**A**) Reference-free 2D class averages generated with limited alignment and reconstruction resolutions to resolve both filaments. Scale bar, 30 nm. (**B**) Reference-free 2D class averages from high-resolution local alignment and corresponding matched projections of paired T-plastin decorated filaments positioned in 3D. Scale bar, 15 nm. (**C**) After initially positioning paired 3D volumes based on front view, cross-correlation between projections of models from side view as one filament is displaced in Z perpendicular to the front view plane and reference-free class averages corresponding to side view. The maximum cross-correlation coefficient is marked with a dashed line. (**D**) Initial 3D models for actin filaments bridged by T-plastin in parallel (left) and anti-parallel (right) configurations. The cross-linking bridge is highlighted in green.

**Fig. S10.**
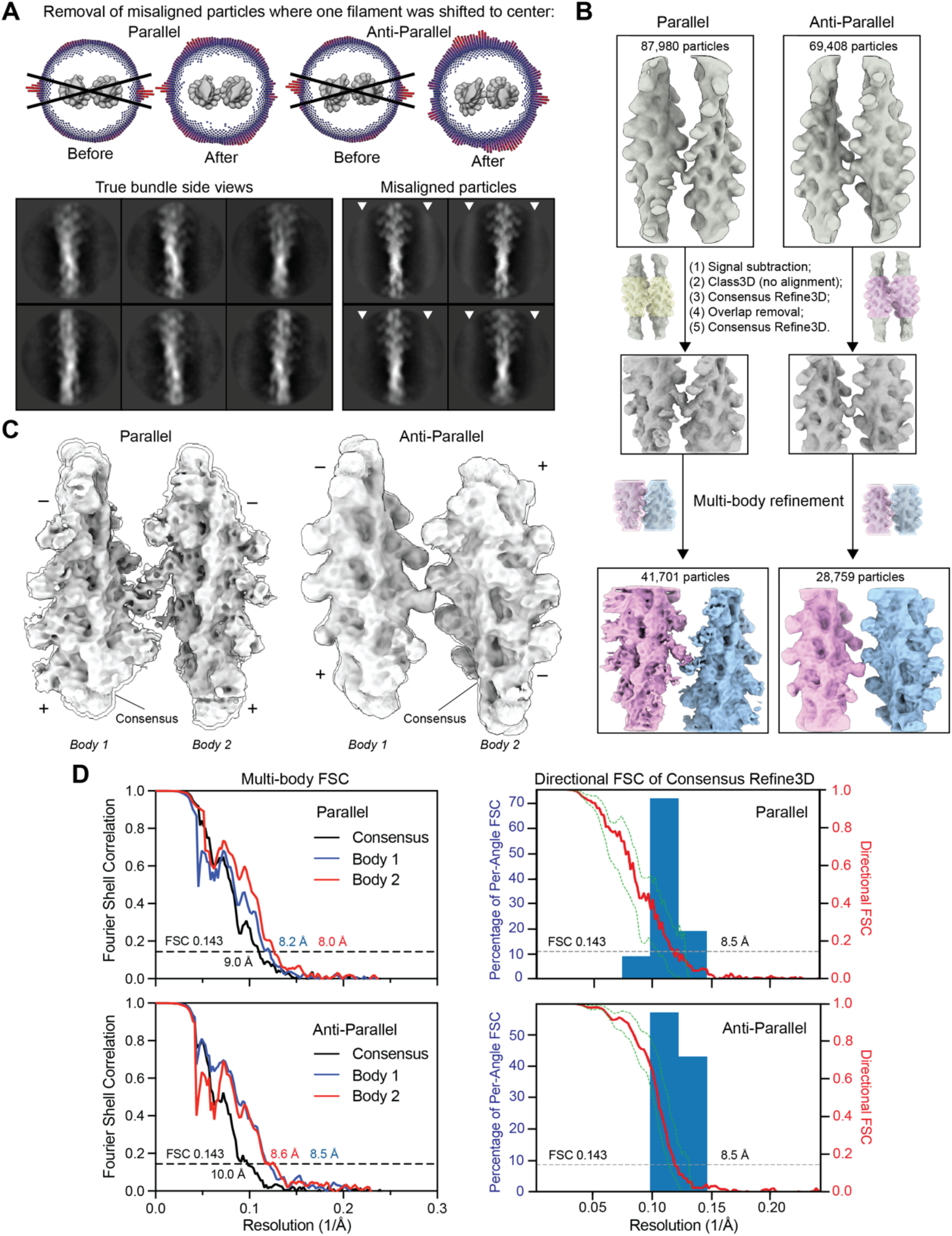
Bundle cryo-EM 3D refinement workflow and resolution assessment. (**A**) 3D angular distributions of particle images prior to removing misaligned particles revealed an over-representation of side views inconsistent with their relatively low frequency in the raw data (top left). 2D classification of particles within the indicated angular wedge without alignment recovered true side views (bottom), as well as views with one filament centered surrounded by a misaligned haze (white arrowheads). Removing these misaligned particles substantially improved the 3D angular distributions (top right). (**B**) Multi-body refinement workflow. (**D**) Resolution assessment of multi-body refinements (left) and directional Fourier Shell Correlation of the consensus Refine3D (right). Dashed green curves represent +/- S.D. from mean of directional FSC.

**Fig. S11:**
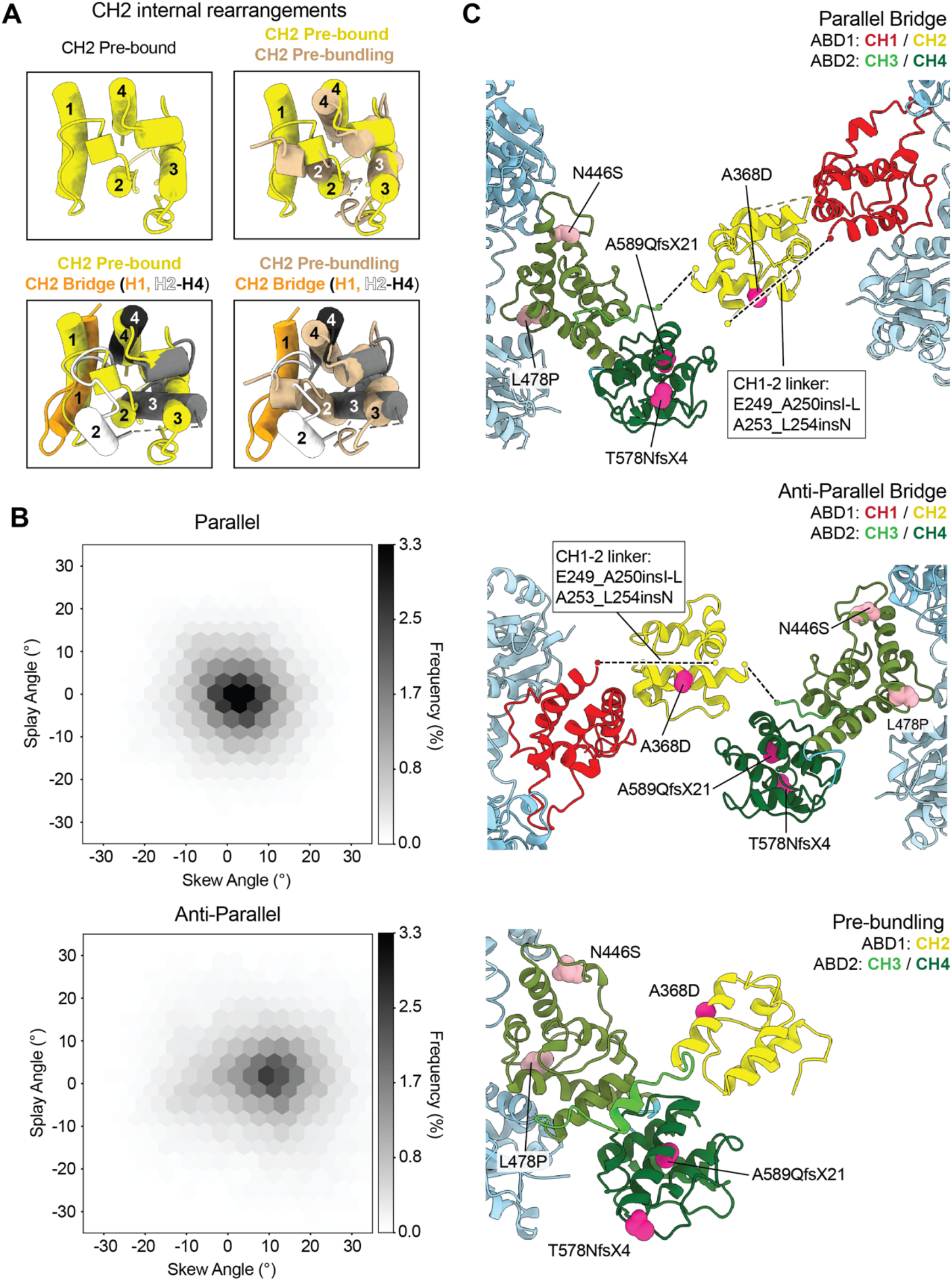
Additional analysis of T-plastin rearrangements and mapping of disease mutations. (**A**) Conformation of the pre-bound T-plastin CH2 domain (top left) and superpositions of the pre-bound, pre-bundling, and parallel bridge conformations of the T-plastin CH2 domain. (**B**) Replotting of measurements from Fig. 3F; there is no apparent coupling between skew and splay angles, suggesting T-plastin acts as a flexible joint within the allowable conformational space of each bridge configuration. (**C**) Osteoporosis-linked T-plastin mutations mapped onto the parallel bridge (top), the anti-parallel bridge (middle), and the pre-bundling (bottom) states of T-plastin.

**Fig. S12.**
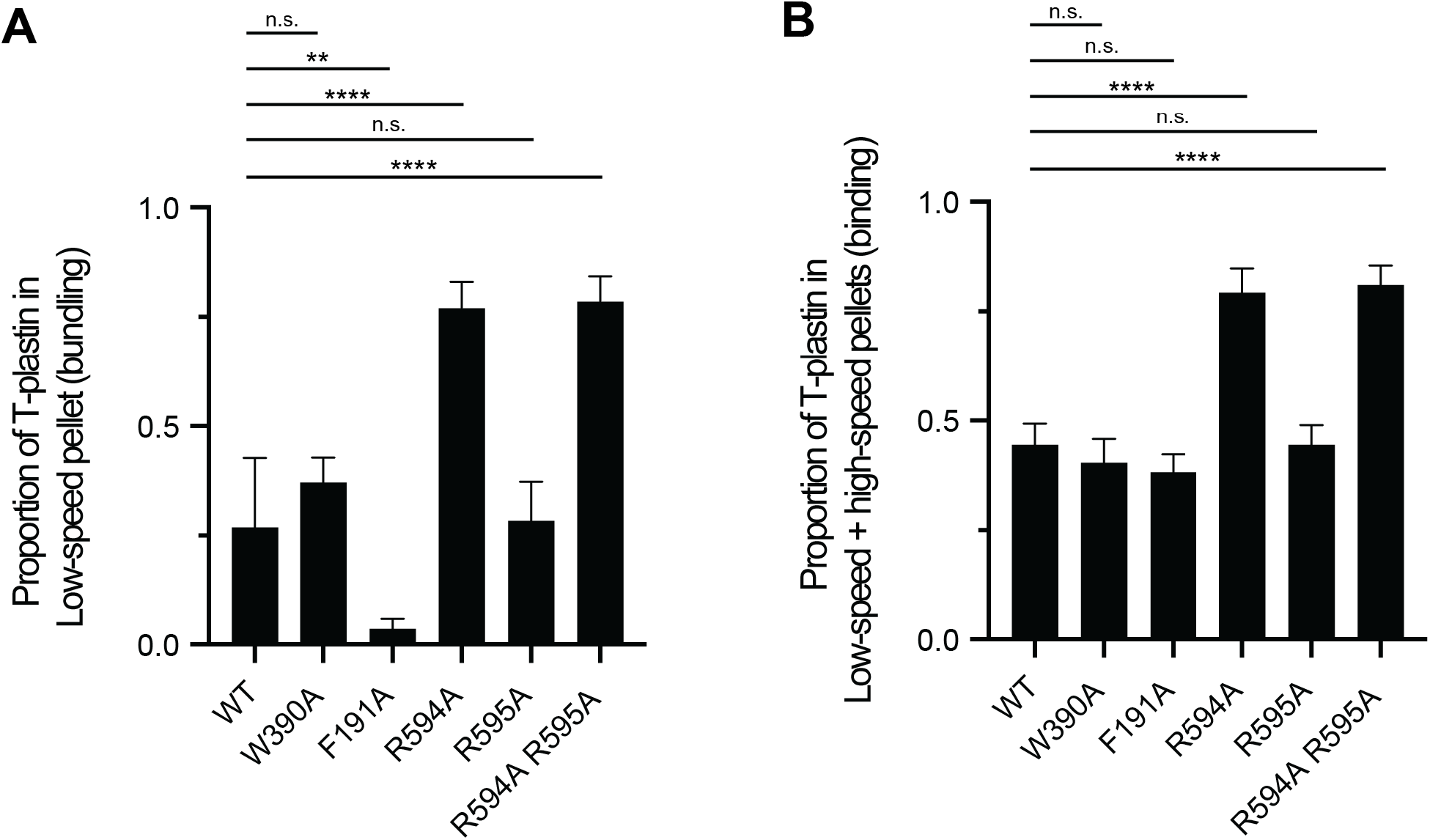
Statistical analysis of co-sedimentation assays. Replotting of data from Fig. 4C **and** Fig. S13D as individual bars for proportion of T-plastin in (**A**) low-speed pellet (WT / W390A: p = 0.16; WT / F191A: p = 0.008; WT / R594A: p = 0.00006; WT / R595A: p = 0.85; WT / R594A R595A: p = 0.00005) and (**B**) low-speed pellet + high-speed pellet (WT / W390A: p = 0.26; WT / F191A: p = 0.06; WT / R594A: p = 0.00006; WT / R595A: p = 0.99; WT / R594A R595A: p = 0.00005). 4 ≤ n ≤ 7. n.s.: not significant, **p < 0.01,****p < 0.0001, T-test.

**Fig. S13.**
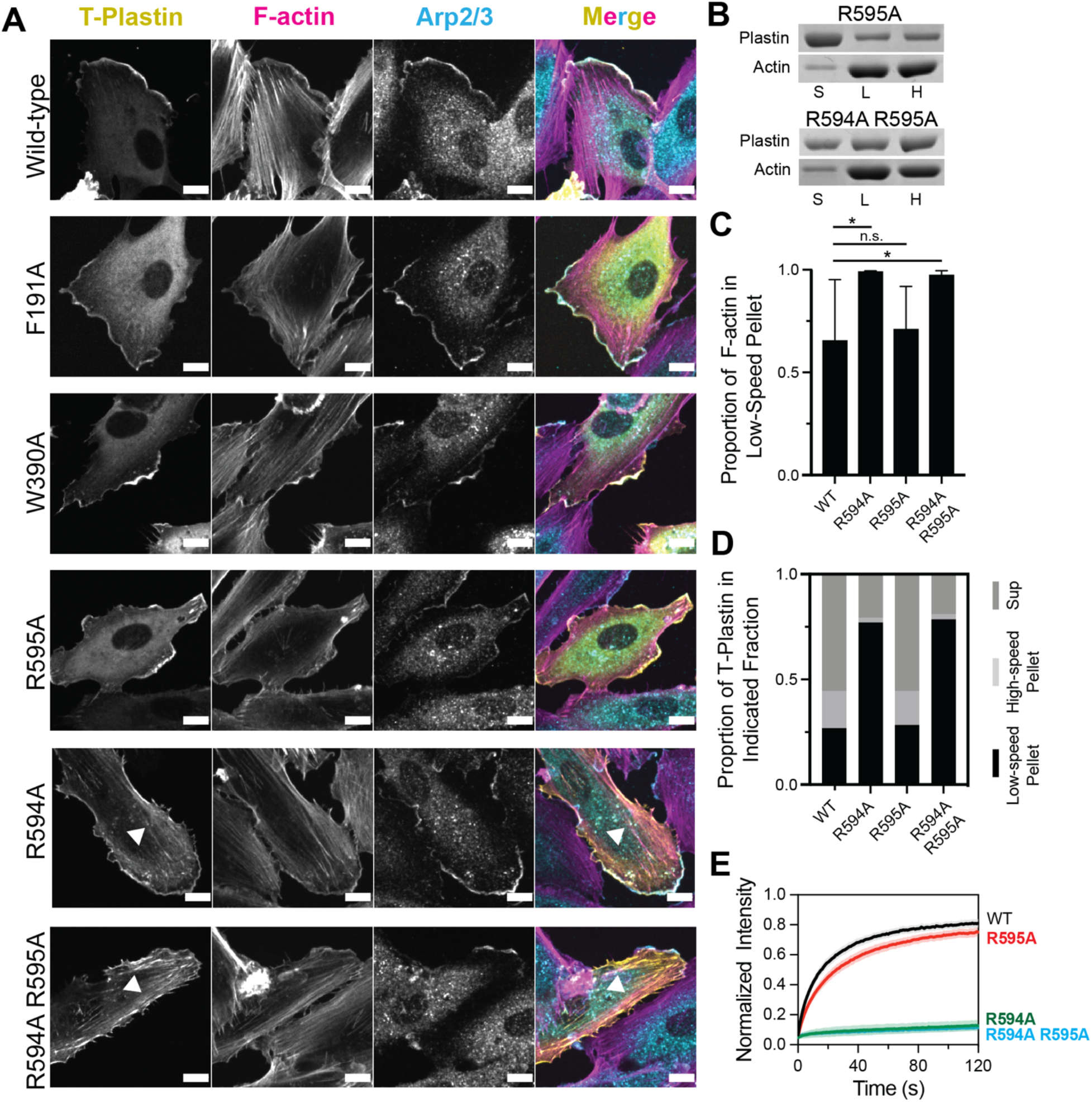
Additional analysis of T-plastin mutants. (**A**) Fluorescence imaging of the indicated T-plastin constructs expressed in HUVEC cells. T-plastin: eGFP; Arp2/3 (marking branched actin at the leading edge): ARP3 immunofluorescence; F-actin: phalloidin. Scale bar, 10 μm. Arrowheads indicate stress fiber / focal adhesion localization in R594A and R594A R595A constructs (bottom 2 rows). (**B**) SDS-PAGE of low-speed / high-speed F-actin co-sedimentation assays with the indicated constructs. S, supernatant; L, low-speed pellet; H, high-speed pellet. (**C**) Quantification of **B**: proportion of F-actin in low-speed pellet (indicative of bundling). Error bars represent S.D., 4 ≤ n ≤ 7. WT / R594A: p = 0.03; WT / R595A: p = 0.74; WT / R594A R595A: p = 0.04. n.s.: not significant, *p < 0.05, T-test. Wild-type and R594A are replotted from Fig. 4C. (**D**) Quantification of **B**: proportions of indicated T-plastin constructs in each fraction. Wild-type and R594A are replotted from Fig. 4D. See Fig. S13 for statistical analysis. (**E)** FRAP assays of the indicated eGFP tagged T-plastin constructs.

**Supplementary Table 1.**
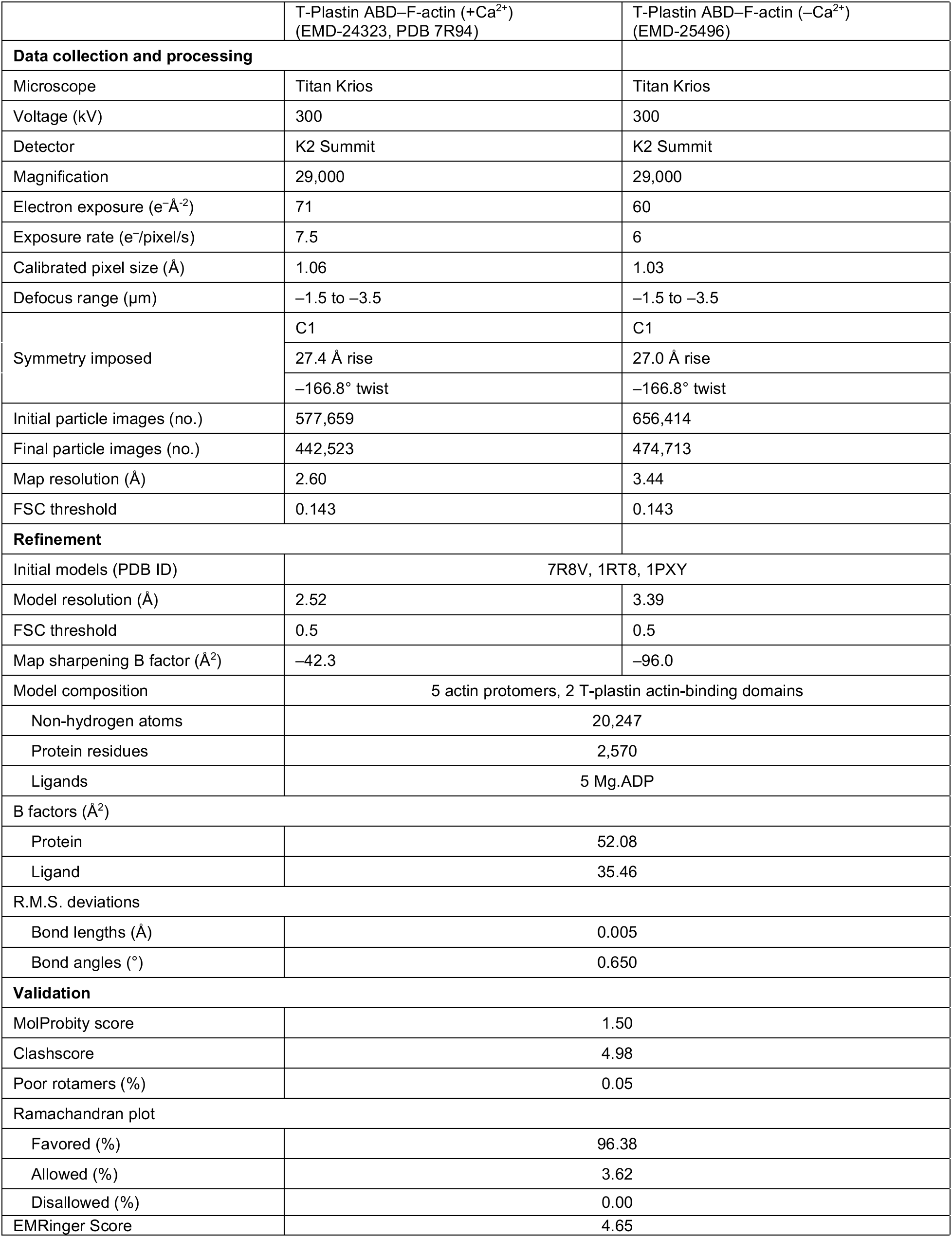
High-resolution cryo-EM data collection, refinement, and validation statistics.

**Supplementary Table 2.** Sub-nanometer cryo-EM data collection, refinement, and validation statistics.

|  | T-Plastin ABD–F-actin,<br>pre-bundling<br>(EMD-25496, PDB 7SXA) | T-Plastin ABD–F-actin,<br>parallel bundle<br>(EMD-25494, PDB 7SX8) | T-Plastin ABD–F-actin,<br>anti-parallel bundle<br>(EMD-25495, PDB 7SX9) |
| --- | --- | --- | --- |
| Data collection and processing |  |  |  |
| Microscope | Titan Krios |  |  |
| Voltage (kV) | 300 |  |  |
| Detector | K2 Summit |  |  |
| Magnification | 29,000 |  |  |
| Electron exposure (e<sup>−</sup>Å<sup>−2</sup>) | 60 |  |  |
| Exposure rate (e<sup>−</sup>/pixel/s) | 6 |  |  |
| Calibrated pixel size (Å) | 1.03 |  |  |
| Defocus range (μm) | −1.5 to −3.5 |  |  |
| Symmetry imposed | C1 |  |  |
| Initial particle images (no.) | 322,743 | 87,980 | 69,408 |
| Final particle images (no.) | 31,413 | 41,701 | 28,759 |
| Map resolution (Å) | 6.85 | 9.04 (consensus)<br>8.16 (body 1)<br>7.95 (body 2) | 9.97 (consensus)<br>8.45 (body 1)<br>8.56 (body 2) |
| FSC threshold | 0.143 |  |  |
| Refinement |  |  |  |
| Initial models (PDB ID) | 7R94 (this work), 1RT8, 1PXY |  |  |
| Map sharpening B factor (Å<sup>2</sup>) | −216.4 | −600 (consensus)<br>−450 (body 1/2) | N.A. (consensus)<br>−300 (body 1/2) |
| Model composition | 3 actin protomers,<br>1 T-plastin | 6 actin protomers,<br>1 T-plastin |  |
| Non-hydrogen atoms | 11,272 | 21,036 | 21,036 |
| Protein residues | 1,431 | 2,668 | 2,668 |
| Ligands | 3 Mg.ADP | 6 Mg.ADP |  |
| R.M.S. deviations |  |  |  |
| Bond lengths (Å) | 0.007 | 0.002 | 0.004 |
| Bond angles (°) | 1.134 | 0.660 | 0.893 |
| Validation |  |  |  |
| MolProbity score | 1.82 | 1.91 | 1.54 |
| Clashscore | 9.77 | 13.07 | 4.78 |
| Poor rotamers (%) | 0.99 | 0.18 | 1.06 |
| Ramachandran plot |  |  |  |
| Favored (%) | 95.55 | 95.89 | 95.97 |
| Allowed (%) | 4.10 | 4.03 | 3.84 |
| Disallowed (%) | 0.35 | 0.08 | 0.19 |

**Supplementary Table 3.** FRAP fitting parameters and analysis.

| **T-plastin construct** | **k (s<sup>-1</sup>)** | **t<sub>1/2</sub> (s)** | **Mobile fraction (%)** | **P value versus wild-type** |
| --- | --- | --- | --- | --- |
| Wild-type | 0.04 | 17.7 | 82 | - |
| F191A | 0.07 | 8.8 | 87 | 0.0001 |
| W390A | 0.02 | 34.6 | 76 | 0.0144 |
| R594A | 0.003 | 212.9 | 25 | < 0.0001 |
| R595A | 0.02 | 27.9 | 80 | 0.0048 |
| R594A R595A | 0.001 | 515.2 | 34 | < 0.0001 |

## Supplementary Movie Captions

**Movie S1. Interpolation along the first principal component from the multi-body refinement of the parallel bundle.** The measured splay and skew angles are displayed at each frame.

**Movie S2. Interpolation along the first principal component from the multi-body refinement of the anti-parallel bundle.** The measured splay and skew angles are displayed at each frame.

**Movie S3. Sequence of T-plastin conformational transitions resulting in parallel bundle.** Colored as in Fig. 3A.

**Movie S4. Sequence of T-plastin conformational transitions resulting in anti-parallel bundle.** Colored as in Fig. 3A.

## Materials and Methods

### Cloning, Expression, and purification of T-plastins

Sequences for the WT, full-length human T-plastin (1–630), as well as T-plastin mutants F191A, W390A, R594A, R595A, and R594 R595A were inserted into a pE-SUMO vector with an N-terminal His6-tag and a SUMO-tag. Sequence of the SUMO protease Ulp1 was inserted into a pET vector with an N-terminal His6-tag. All T-plastin constructs and the and SUMO protease Ulp1 were expressed in Rosetta2(DE3) E. coli cells (Novagen) grown in LB media at 37°C to an optical density of 0.8-1.0 and induced with 0.7 mM IPTG. After induction, the cells were grown for 16 hours at 16°C, then cell pellets were collected and stored at −80°C until use. Cell pellets were resuspended in Lysis Buffer (50 mM Tris-Cl pH 8.0, 150 mM NaCl, 5% v/v glycerol, 2 mM β-mercaptoethanol, 20 mM imidazole) and lysed with an Avestin Emulsiflex C5 homogenizer, after which the lysate was clarified at 15,000 g for 30 minutes. Cleared lysate was incubated with Ni-NTA resin (Qiagen) for 1 hour on a rotator at 4°C, after which the flow-through was discarded and the resin was washed with 5 bed volumes of lysis buffer. Proteins were subsequently eluted in Elution Buffer (50 mM Tris-Cl pH 8.0, 150 mM NaCl, 5% v/v glycerol, 2 mM β-mercaptoethanol, 300 mM imidazole). Purified His-tagged SUMO protease Ulp1 was then added at 0.05 mg / ml working concentration into the protein solutions of SUMO-tagged proteins, then the protein solution was dialyzed against Dialysis Buffer (20 mM Tris-Cl pH 8.0, 300 mM NaCl, 5% v/v glycerol, 2 mM β-mercaptoethanol) for 16 hours. The protein solution was then reapplied to Ni-NTA resin, and the flow-through was collected. Protein was then sequentially purified by a HiTrapQ HP anion exchange column (GE Healthcare) followed by size exclusion chromatography on a Superdex 200 Increase column in Gel Filtration Buffer supplemented with 10% v/v glycerol, then snap-frozen in liquid Nitrogen and stored at −80°C.

### Cryo-EM sample preparation and data collection

Filamentous actin (F-actin) was polymerized in G-Mg (G buffer supplemented with 0.1 mM MgCl_2_) and KMEI (50 mM KCl, 1 mM MgCl_2_, 1 mM EGTA, 10 mM imidazole pH 7.0, 1 mM DTT) buffers as described previously (*1*) from 5 µM globular actin (G-actin) monomers at room temperature for 1 hr, then diluted to 0.6 μM in KMEI prior to use. Full-length human T-plastin was buffer exchanged into KMEI or Ca-KMEI (KMEI buffer + 2mM CaCl_2_) by desalting column (GE PD SpinTrap G-25) and was maintained at a concentration of 20 μM. All KMEI buffers were supplemented with 1 mM DTT and 0.05% NP-40.

Immediately prior to sample preparation, CF-1.2/1.3-3Au 300-mesh gold C-flat holey carbon cryo-TEM grids (Protochips) were plasma cleaned with a Hydrogen / Oxygen mixture for 5 seconds in a Gatan Solarus. Actin solution (3 μl) was first applied to the grid in the humidified chamber of a Leica EM GP plunge freezer and incubated for 60 s at 25°C. T-plastin solution (3 μl) was then applied and incubated for 30 s. Solution (3 μl) was then removed and an additional 3 μl of T-plastin solution was applied. After an additional 30 s, the grid was back-blotted for 5 s, plunge-frozen in ethane slush, and stored in liquid Nitrogen until imaging.

Cryo-EM data for the T-plastin–actin (–Ca^2+^) complex were recorded on a Titan Krios (ThermoFisher / FEI) at the Rockefeller University operated at 300 kV equipped with a Gatan K2 Summit camera. SerialEM (*2*) was used for automated data collection. Movies were collected at a nominal magnification of 29,000X in super-resolution mode resulting in a calibrated pixel size of 1.03 Å / pixel (super-resolution pixel size of 0.515 Å / pixel), over a defocus range of −1.5 to −3.5 μm. 40 frames were recorded over 10 s of exposure at a dose rate of 6 electrons per pixel per second (1.5 electrons per Å^2^ per second) for a cumulative dose of 60 electrons per Å^2^. The T-plastin–actin (+Ca^2+^) dataset was recorded at the New York Structural Biology Center (NYSBC) on a Titan Krios operated at 300 kV also equipped with a Gatan K2 Summit camera. Movies were collected in counting mode resulting in a calibrated pixel size of 1.06 Å / pixel, over a defocus range of −1.5 to −3.5 μm. 50 frames were recorded over 10 s of exposure at a dose rate of 1.34 electrons per pixel per second (1.4204 electrons per Å^2^ per second) for a cumulative dose of 71.02 electrons per Å^2^.

### High-resolution cryo-EM image processing

Unless otherwise noted, all image processing was performed within the RELION-3.0 package (*3*), following a recently described procedure (*4*). Movie frames were aligned and summed with 2 x 2 binning for the –Ca^2+^ dataset, and no binning for the +Ca^2+^ dataset, using the MotionCor2 algorithm (*5*) implemented in RELION (*6*) with 5 × 5 patches. The contrast transfer function (CTF) was estimated from non-doseweighted summed images with CTFFIND4 (*7*). Bimodal angular searches with psi angle priors were used in all subsequent 2D and 3D alignment / classification procedures. Approximately 2,000 segments were initially manually picked, extracted, and subjected to 2D classification to generate templates for auto-picking. Helical auto-picking was then performed with a step-size of 3 asymmetric units corresponding to a 27 Å helical rise. Segments were extracted from dose-weighted (*8*) sum images in 512 x 512 pixel boxes without down-sampling, followed by a second round of 2D classification and auto-picking with featureful class averages. Auto-picked segments were then extracted and subjected to 2D classification using a 200 Å tube diameter and 300 Å mask diameter. Segments that contributed to featureful class averages were selected for 3D analysis.

All subsequent 3D analysis steps were primed with estimates of helical rise and twist of 27.0 Å and −167.0°, respectively, utilizing an initial reference low-pass filtered to 35 Å resolution, with the outer tube diameter set to 200 Å, inner tube diameter set to −1, and the mask diameter set to 300 Å. The first round of 3D classification into 3 classes was performed with bare actin filament reconstruction (EMBD-7115) as the initial reference. A second iteration was then performed with a class featuring clear ABP density from first round as the initial reference. For the T-plastin–actin (–Ca^2+^) dataset, this second round of 3D classification yielded two classes with helical parameters similar to the initial estimates and well-resolved 3D features, and one junk class with aberrant helical parameters and distorted features. For T-plastin–actin (+Ca^2+^) dataset, all three classes were good, featureful 3D classes, indicating the high quality of this dataset. Segments contributing to the selected classes were then pooled for 3D auto-refinement.

The first round of auto-refinement was then performed using one good 3D class as the initial reference. All masks for subsequent post-processing steps were calculated with 0 pixel extension and a 6 pixel soft edge from the corresponding converged reconstruction, low-pass filtered to 15 Å and thresholded to fully encompass the density. First-round post-processing was performed with a 50 % z length mask, followed by CTF refinement without beam-tilt estimation and Bayesian polishing (*6*). A second round of auto-refinement was then performed, followed by post-processing with a 30 % z length mask, then a second round of CTF refinement with beam-tilt estimation and Bayesian polishing. Final auto-refinement was then performed, once again employing a 30 % z length mask for post processing.

The final reconstructions converged with helical rise of 27.0 Å and twist of −166.8° for the T-plastin–F-actin ABD-F-actin complex (–Ca^2+^), and a helical rise of 27.4 Å and twist of −166.8° for the T-plastin–F-actin ABD-F-actin complex (+Ca^2+^), consistent with our finding that actin rearrangements evoked by T-plastin are minimal (Fig. S2C). Global resolution estimates were 3.4 Å for the full-length T-plastin–F-actin complex (–Ca^2+^), and 2.6 Å for the full-length T-plastin–F-actin complex (+Ca^2+^), by the gold-standard Fourier shell correlation (FSC) 0.143 criterion. The B-factors estimated during post-processing were then used to generate sharpened, local-resolution filtered maps with RELION. Key statistics summarizing high-resolution cryo-EM image processing are reported in Table S1.

### T-plastin pre-bundling state cryo-EM image processing

The weak signal in Class I of the –Ca2+ condition (Fig. S1) suggested that in a subset of the particle images, signal from an additional CH domain was consistently present in a defined region. To isolate and refine those particles, symmetry expansion and subsequent rounds of focused 3D classification were used to generate a set of 322,743 segment images (Fig. S5A). Using the asymmetric unit and helical parameters from 3D auto-refinement, particles were symmetry expanded. These particles were then re-extracted with recentering of 2D shifts with a box size of 128 pixels at a down-sampled pixel size of 4.12 Å and subjected to a consensus 3D auto-refinement. After this consensus refinement, a mask containing only density for the bound ABD and the additional density was used for an initial round of 3D classification with 14 classes and no image alignment. Classes with consistent density were selected and subjected to 3D auto-refinement. The resultant map was used as an updated reference for another round of 3D classification of the original 322,743 segment images with 6 classes, using an updated mask. After selecting 147,851 segment images from two classes with similarly placed density, a local 3D auto-refinement was performed, which showed substantially increased density in the masked region beyond the bound ABD. Using an updated mask, another round of 3D classification was run with 3 classes and no image alignment. One class, composed of 50,077 particles, with good density was selected. These particles were re-extracted at a box size of 384 pixels without down-sampling and locally refined using a 70% mask. These particles were then re-extracted with the plastin molecule at the center of the box and re-refined with a 50% mask. A subsequent round of 3D classification with 3 classes and no image alignment and was performed, and the best two classes were selected with 31,413 particles. After a local 3D auto-refinement, the particles were re-centered on the actin filament to optimize CTF refinement and Bayesian Polishing performance for two rounds. During the second round of particle polishing, the particles were once again re-centered on the bound T-plastin protein. Polished particles were subjected to a final 3D auto-refinement, which yielded a map assessed at 4.4 Å resolution. A final, local 3D auto-refinement was performed by masking only the plastin and two actin protomers to give the final map at 6.9 Å resolution (Fig. S5) within this masked region.

### Synthetic dataset generation

The high similarity between projection images of individual actin filaments and two-filament actin bundles prevents simple, cross-correlation-based picking from distinguishing between bundled and unbundled actin. We developed a more sophisticated convolutional neural network-based approach to specifically identify bundled filaments, while excluding individual filaments. To achieve full-micrograph segmentation, a neural network was first trained to recognize potential bundle configurations in synthetic data, and then used to predict on real data.

We first trained a denoising autoencoder on projections of plausible *in silico* bundle models and used the learned weights from this network to make a semantic segmentation network. The precise workflow for synthetic projection generation is outlined in Fig. S6. Briefly, projection images were generated of zero to three filament units, with each filament unit consisting of either an individual filament or a two-filament bundle. To approximately reflect the frequency of bundles in the actual dataset, each filament unit had a 65% chance of being a bundle and 35% chance of being an individual filament. If the filament unit was a bundle, the filament would be copied, and the copy would be rotated about its helical axis by a random, uniformly sampled integer between 0° and 359°, randomly tilted by a bimodal Gaussian centered at 0° and 90°, with standard deviations of 1.5°, and then randomly translated in the y-direction by 159 Å ± 24 Å (empirically measured from 127 bundles in real micrographs), and uniformly translated in the z-direction (along the helical axis) by ±181 Å. Subsequently, for both individual and bundled filament units, the filament unit would be rotated about the phi and rot angles by a random, uniformly sampled value between 0° and 359°, and the tilt by 0° with a standard deviation of 2.5°. The filament unit was then randomly translated around the box by ±250 Å and projected along the z-axis to generate a noiseless projection. The same map was used to generate a noisy image paired with this noiseless projection, by adding pink noise in Fourier space, as implemented in EMAN2’s python package to generate realistic-looking synthetic data (*9*). Three-channel stacks of semantic maps associated with the noisy/noiseless projection pairs were generated by binarizing the filament unit and assigning it as a bundle or individual filament before projection.

### Network architecture and training

A denoising autoencoder (DAE) was trained using the architecture outlined in Fig. S7A. Each trainable layer had a ReLU activation function, except the final layer which had a linear activation function. The negative of the cross-correlation coefficient was used as the loss function. For training, the weights were initialized using the default initialization in TensorFlow. The model was trained using the Adam optimizer version of stochastic gradient descent with a learning rate of 0.00005 and minibatch size of 16 until the model converged (no improvement in validation loss for 3 epochs). Upon network convergence, the weights from the best epoch were restored. For training, 300,000 noisy / noiseless projection pairs with box sizes of 192×192 were generated,90% of which was used for training and 10% for validation. Upon network convergence, the denoising autoencoder had an average cross-correlation coefficient of 0.984 on the validation set.

After training the model as a DAE, a semantic segmentation network was trained by copying all layers and weights of the DAE, except for the final layer, which was replaced with a 3-channel layer with softmax activation and default TensorFlow initialization. This semantic segmentation network was then trained with a learning rate of 0.00001. For training, 50,000 pairs of noisy inputs and semantically segmented targets of dimension 192×192 and 192×192×3, respectively, were used with a minibatch size of 16; 90% of the synthetic data was used for training and 10% for validation. The loss function was categorical cross-entropy, and upon network convergence the model had a categorical cross-entropy of 0.069 on the validation set. Example network performance on synthetic data is shown in Fig. S7A,B.

Models were trained on a single NVIDIA 2080 Ti GPU with 11 GB of VRAM. Training required approximately 2.25 h per epoch for the denoising autoencoder and 0.3 h per epoch for the semantic segmentation network. Validation loss began to plateau around the fifth epoch for both models and then slowly improved until convergence (Fig. S7C). As a separate estimate of the 2D denoising reconstruction’s resolution, 10,000 noiseless synthetic particle images not used during network training or validation were compared to noisy particle images denoised using the trained DAE, and the Fourier Ring Correlation (FRC) was computed (Fig. S7D); the average FRC curve fell below 0.5 at 12.2 Å (∼1.5 times Nyquist resolution), and 100 example FRC curves are also shown.

### Particle picking using neural network

To pick particles, motion-corrected micrographs down-sampled by 4 were converted into semantic maps by extracting 192-pixel boxes across the micrograph in a raster pattern with 48 pixels of overlap and stitching back the output into a semantic map by computing a maximum intensity projection of the overlapping regions. Only the bundle channel results were used, and they were binarized using a fixed threshold of 0.85. After binarization, central axes of the bundles were selected for by excluding pixels near object borders using empirically derived parameters, and small objects were removed. The remaining binarized image was skeletonized, and non-maximum suppression was used to ensure all particle picks were spaced at least 148 Å away from each other. These particle coordinates were used for extraction in RELION.

### Bundle processing

Visual inspection revealed that nearly all extracted particles had multiple filaments, but reference-free 2D classification with standard parameters produced classes with one filament or one well-resolved filament and one poorly resolved filament (not shown). To prevent alignment from refining one filament at the expense of the other, particles images were extracted in a large, 256-pixel box downsampled to 4.12 Å pixel size, and multiple rounds of 2D classification were performed in cryoSPARC (*10*), limiting the reconstruction resolution to 45 Å and the alignment resolution to 50 Å (Fig. S9A); the “Align filament classes vertically” option was used to determine in-plane rotation, and at each iteration, 2D class averages were re-centered using a binary mask with a low threshold to maintain the bundle in the class average center. Particles were re-extracted with shifted, re-centered coordinates and psi angles were rotated by 90° for RELION helical conventions. With these re-centered particles, 2D classification with small translations yielded high-quality, reference-free 2D class averages in RELION, where obvious parallel and anti-parallel 2D classes were present (Fig. S9B). Despite exhaustive attempts at generating reasonable initial models using *ab initio* model generation, models would frequently be produced with one filament centered in the box or two poorly defined filaments. Therefore, using the reference-free 2D class averages, initial models were generated using a custom projection-matching scheme. The map derived from single-filament helical analysis was rescaled and roughly positioned in the box to project onto one of the two filaments, then EMAN2’s e2classvsproj script (*9*) was used on each filament to globally search Euler angles (with 10 degree sampling) and shifts (maximum of 84 A) for an initial projection-match to the 2D class average. Finally, a custom projection-matching script using the EMAN2 python package was used to perform a finer, gradient-descent-based projection matching of each filament. While these projections had excellent correspondence to the 2D class averages, they did not have full 3D information to properly position filaments for an initial model. To properly z-position the filaments, a parallax-based approach using 2D classes that had clear side-views was used, and adjustments to the relative z-positions of the oriented filament maps was performed to maximize the cross-correlation between the side-view 2D class averages and the 3D models (Fig. S9C). These maps were then lowpass filtered to 20 Å and used as initial references for 3D classification (Fig. S9D).

3D classification required careful angular and translational searches to prevent head-on two-filament images from shifting to side-on views of one filament. An approach similar to previous 3D classification schemes of large, mixed-population filamentous structures was used (*11*). Briefly, the tilt prior was kept fixed at 90° and global searches of rot, local, bimodal searches of psi, and local searches of tilt with 3.7° sampling and a 20° search range were performed, with a translation search range of 41.2 Å. After one iteration with these parameters, the 3D classification was interrupted; the tilt prior was updated, and translations were deleted. The 3D classification then resumed for three iterations, with global searches of rot, local, bimodal searches of psi, and local searches of tilt with a fine 1.8° sampling and 5° search range, and a large 123.6 Å translational search range was used. During this supervised 3D classification, there was an apparent preferred orientation for particular rot angles (Fig. S10A). 2D classification of particles with rot angles of 0±15° or 180±15° revealed that some side-on view 2D class averages look similar to individual filaments, and their constituent particle images were either bundled particles shifted to have one filament in the box center or poorly picked individual filaments. 2D classes with one centered filament that was much better resolved than other 2D classes were excluded and the angular distribution improved (Fig. S10A, B). After this winnowing, five more iterations of 3D classification were performed with a smaller translational search of 20.6 Å, leading to 87,980 particles sorted into the parallel class and 69,408 particles sorted into the anti-parallel class (Supplementary Fig.10C). A 45% z-mask was generated for both classes, and subsequent signal subtraction and focused 3D classification without alignment removed particles without clear bridging plastin density. Un-subtracted particles were re-substituted, and following a consensus 3D auto-refinement, particles overlapping within the masked region were removed. A final, asymmetric, consensus 3D auto-refinement was performed on 41,701 particles for the parallel class, which reached 9.0 Å, and 28,759 particles for the anti-parallel class, which was assessed at 10.0 Å (Supplementary Fig. 10D,E).

### Variability analysis

During 3D auto-refinement, it became apparent that filament density quickly smeared outside the masked region, presumably due to the relative flexibility of the complex. This observation, coupled with the composition of the system being two rigid bodies connected by a flexible cross-linking protein, led us to employ multi-body refinement, as implemented in RELION, to handle flexural heterogeneity (*12*). Using default 3D multi-body refinement parameters, with masks shown in Fig. S10C, the resolution of the constituent filaments improved. Furthermore, we reasoned we could utilize the multi-body refinement parameters to measure the relative motions of the two filaments to each other. Specifically, volumes were generated for each particle image using RELION’s relion_flex_analyze script (41,701 volumes for parallel, and 28,759 volumes for anti-parallel), and atomic models of two separate plastin-decorated actin filaments were procedurally rigid-body docked into each of the volumes using scripting functions in UCSF Chimera (*13*). Distances and measurements between these docked models were measured using custom scripts employing functions from the ProDy package (*14*). Plots were generated with GraphPad Prism (Fig. 3E-H).

### Model building and refinement

To generate homology models of the actin-binding ‘core’ (Fig. 1B) of plastin, the Robetta server was used (*15*). The selected homology model for ‘pre-bound’ T-plastin was the model that had the highest score. Sharpened, local-resolution-filtered maps as described above were used for model building. The high-resolution density maps were of sufficient quality for *de novo* atomic model building. As structures of components were available, initial models of actin (PDB 3j8a) and the ‘pre-bound’ T-plastin homology model were fit into the density map using Rosetta (*16*). Models were subsequently inspected and adjusted with Coot (*17, 18*), and regions that underwent significant conformational rearrangements were manually rebuilt. The models were then subjected to several rounds of simulated annealing followed by real-space refinement in Phenix (*19, 20*) alternating with manual adjustment in Coot. A final round of real-space refinement was performed without simulated annealing.

The pseudo-atomic models for the T-plastin pre-bundling state (Fig. 2) and both T-plastin bundle configurations (Fig. 3) were generated by rigid-body docking the high-resolution post-bound T-plastin model and ABD1 fragments from the ‘pre-bound’ homology model, followed by flexible fitting with ISOLDE (*21*), then real-space refinement in Phenix with harmonic restraints on the starting model enabled. Key statistics summarizing model building, refinement, and validation are reported in Table S1.

Structural figures and movies were prepared with ChimeraX (*22*). Per-residue RMSD analysis was performed with UCSF Chimera (*13*) as previously described (*23*). The surface area of actin-binding interfaces was calculated with PDBePISA (*24*) (EMBL-EBI). Model quality was assessed with EMRinger (*25*) and MolProbity (*26*) as implemented in Phenix.

### Sequence alignments

Protein sequences of human I-plastin (Q14651), human L-plastin (P13796), human T-plastin (P13797) were aligned with ClustalOmega (*27*) (EMBL-EBI).

### Actin co-sedimentation assays

Briefly, mixtures of F-actin (5 μM) and the indicated T-plastin constructs (2 μM) were incubated at room temperature for 30 min in co-sedimentation buffer (10 mM Tris pH 7.5, 100 mM KCl, 2.5 mM MgCl_2_, and 2 mM DTT. The samples were then spun for 30 min. at 16,000 rpm (low-speed) in a TLA-100 rotor and polycarbonate centrifugation tubes (Beckman Coulter No.343775). This pellet was the ‘low-speed’ pellet. The supernatant was taken out from the centrifugation tube and then spun in a fresh centrifugation tube for another 30 min, at 100,000 rpm (high-speed). This pellet was the ‘high-speed’ pellet, and the supernatant was also collected. All three fractions were subject to SDS-PAGE and Coomassie staining. The gels were scanned using LI-COR imaging system, and subsequently quantified with ImageJ. Plots were generated with GraphPad Prism, and statistical analysis was performed with Microsoft Excel.

### Cell culture and transfections

All experiments using HUVEC were performed with an hTERT-immortalized HUVEC line previously described (*28*) and cultured in EGM-2 media (Lonza, CC-3162). For transient DNA transfections, 1.5 × 10^4^ HUVE, were plated the day before transfection in a glass-bottom 96-well plate (Cellvis) coated with 31 µg/mL collagen (Advanced BioMatrix, 5005-100 ML). The day of transfection, culture medium was replaced with 80 µL of antibiotics-free culture media per well. Then, 0.2 µg DNA encoding GFP-tagged T-Plastin (WT or mutant) and 0.25 µL Lipofectamine2000 (Thermo Fisher Scientific), mixed in 20 µL OptiMEM (Thermo Fisher Scientific), was added following the manufacturer’s instructions. This transfection mix was replaced after 3 h with culture media and cells were processed for imaging after 24 h.

### Fixed fluorescence imaging of cells

For fixed imaging, cells were rapidly fixed in 37 °C 4% paraformaldehyde in PBS, washed in PBS, permeabilized in 0.2% TritonX-100 in PBS, washed again and blocked in 3% FBS in PBS prior to primary and secondary antibody staining. Phalloidin conjugated to Alexa Fluor 568 (A12380, used at 1:500) and a secondary antibody against mouse IgG conjugated to Alexa Fluor 647 (A21235, used at 1:500) were from Thermo Fisher Scientific. The mouse monoclonal antibody against Arp3 (A5979, used at 1:250) was from Sigma.

All images were acquired on an automated confocal system controlled by Slidebook software (Intelligent Imaging Innovations, 3i). The system consists of an Eclipse-Ti body with perfect focus system (Nikon Instruments), CSU-W1 spinning disc (Yokogawa), a 3i laser stack with 405, 445, 488, 515, 561, and 640 nm lines, an environmental chamber (Haison), a 60X 1.27 NA water immersion objective, and two Zyla 4.2 sCMOS cameras (Andor) with motorized dichroics enabling simultaneous acquisition of channels.

### FRAP assays and quantification

For live imaging, cell culture media was replaced with extracellular buffer (ECB) consisting of 125 mM NaCl, 5 mM KCl, 1.5 mM MgCl_2_, 1.5 mM CaCl_2_, 20 mM HEPES, supplemented with 10 mM D-glucose, 1% FBS, and 5 ng/mL bFGF (R&D Systems, 223-FB). To avoid interference from lamellipodial protrusion/retraction cycles on T-Plastin recovery measurements, HUVEC were first treated with 20 µM Y-27632 for 20 min., then a cocktail of 20 µM Y-27632, 8 µM Jasplakinolide, and 5 µM Latrunculin-B (JLY) for 30 min. to freeze actin turnover (*29*). Regions representing protrusions containing GFP-tagged T-plastin were selected for photobleaching with high intensity 488 nm laser light (Vector, Intelligent Imaging Innovations) and imaged every second for at least 3 minutes.

Analysis regions were subsequently defined in Slidebook to exclude the ends of actin stress fibers and focal adhesions, and the region intensities were measured to quantify their recovery after photobleaching. Intensities were background subtracted and normalized to their pre-bleach intensity, then analyzed in GraphPad Prism by fitting to an exponential one-phase association model, Y = Y_0_ + (Plateau - Y_0_) * (1 - e^(-K * x)^). Wild-type and mutant data were compared using a two-way ANOVA with a mixed-effects model and Geisser-Greenhouse correction (Table S3).

